# Continuous self-repair protects vimentin intermediate filaments from fragmentation

**DOI:** 10.1101/2024.09.02.610785

**Authors:** Quang D. Tran, Martin Lenz, Guillaume Lamour, Lilian Paty, Maritzaida Varela-Salgado, Clément Campillo, Hugo Wioland, Antoine Jegou, Guillaume Romet-Lemonne, Cécile Leduc

## Abstract

Intermediate filaments are key regulators of cell mechanics. Vimentin, a type of intermediate filament expressed in mesenchymal cells and involved in migration, forms a dense network in the cytoplasm that is constantly remodeling through filament transport, elongation/shortening, and subunit exchange. While it is known that filament elongation involves end-to-end annealing, the reverse process of filament shortening by fragmentation remains unclear. Here, we use a combination of *in vitro* reconstitution, probed by fluorescence imaging and AFM, with theoretical modeling to uncover the molecular mechanism involved in filament breakage. We first show that vimentin filaments are composed of two populations of subunits, half of which are exchangeable and half immobile. We also show that the exchangeable subunits are tetramers. Furthermore, we reveal a mechanism of continuous filament self-repair, where a soluble pool of vimentin tetramers in equilibrium with the filaments is essential to maintain filament integrity. Filaments break due to local fluctuations in the number of tetramers per cross-section, induced by the constant subunit exchange. We determine that a filament tends to break if approximately four tetramers are removed from the same filament cross-section. Finally, we analyze the dynamics of association/dissociation and fragmentation to estimate the binding energy of a tetramer to a complete versus a partially disassembled filament. Our results provide a comprehensive description of vimentin turnover and reveal the link between subunit exchange and fragmentation.

**SIGNIFICANCE STATEMENT:** Intermediate filaments such as vimentin are key contributors to cell mechanics. These filaments are continuously assembled and disassembled by the cell as it grows and moves. While the assembly process is relatively well characterized, here we investigate their much less understood disassembly. We find that the filaments continuously exchange their constitutive proteins with the surrounding solution throughout their length and when too many subunits are removed, the filament breaks. This process sharply differs from the length regulation mechanisms known to occur in other types of filaments involved in the regulation of cell mechanics. Importantly, we also identify two distinct populations of subunits within the filaments, which raises new questions about vimentin intermediate filament structure.

## INTRODUCTION

Intermediate filaments (IFs) are one of the three major components of the cytoskeleton, along with actin filaments and microtubules, and play a critical role in determining cell shape and mechanics. Notably, IFs contribute to preserving cell integrity by enabling cells to withstand significant deformations (Latorre *et al*. 2018, Hu *et al*. 2019, Nagle *et al*. 2022). Beyond their well-established mechanical roles, IFs are also implicated in various cellular functions, including signaling, division, polarity, and migration (Patteson *et al*. 2020), often through their interaction with the other types of cytoskeletal filaments (Huber *et al*. 2015, Leduc and Etienne-Manneville 2015). Mutations in IF proteins, which can disrupt IF assembly or remodeling, have been associated with numerous pathologies (Omary et al. 2009). However, despite their critical roles, the mechanisms underlying IF assembly and disassembly are much less understood than those of F-actin and microtubules.

IFs constitute a large and heterogeneous family categorized into six main classes. Vimentin, a class III IF, is predominantly expressed in cells of mesenchymal origin and is commonly used as a marker of the epithelial-mesenchymal transition (Kalluri and Weinberg 2009). Vimentin filaments form a dense, interconnected network extending from the nucleus to the plasma membrane, and with dynamic remodeling essential to their cellular functions (Leduc and Etienne-Manneville 2017, Etienne-Manneville 2018, Nunes Vicente et al. 2022). In cells, filament assembly occurs via end-to-end annealing (Çolakoğlu and Brown 2009) while filament disassembly is regulated by post-translational modifications (PTMs) such as phosphorylation (Inagaki et al. 1987, Eriksson et al. 2004, Snider and Omary 2014). Indeed, phosphorylation of vimentin reduces the positive charges in its head domain, thus destabilizing internal interactions and leading to filament disassembly (Eriksson et al. 2004, Kraxner et al. 2021). *In vitro* reconstitution of vimentin filaments with purified proteins has further elucidated the mechanisms of filament assembly (Herrmann et al. 1996, Sokolova et al. 2006, Brennich et al. 2011, Winheim et al. 2011, Nöding et al. 2014, Lopez *et al*. 2016, Schween et al. 2022). These studies demonstrated that assembly initiates with the lateral association of apolar tetramers to form short filament precursors referred to as unit-length filaments (ULFs). ULFs subsequently undergo end-to-end annealing for longitudinal elongation (Herrmann and Aebi 2016). Unlike actin and microtubules, vimentin does not polymerize through subunit addition at filament ends. Once assembled, vimentin filaments exhibit remarkable stability and can remain intact at room temperature for over a week (Fig. S1A)(Herrmann and Aebi 2016). The main way to induce spontaneous disassembly, is to greatly decrease the ionic strength (*<*20 mM) or add denaturing reagents (Herrmann and Aebi 2016).

Although vimentin filaments are highly stable, we have recently demonstrated that they can fragment in the absence of severing proteins or PTMs (Tran et al. 2023). We estimated the mean bond breaking time between successive ULFs to be ~18 h, which is of the same order of magnitude as the filament breaking time observed in cells (Hookway *et al*. 2015). However, the molecular mechanism involved in this filament breakage is unknown. We propose that the local dissociation of subunits from the lattice weakens the filament, eventually leading to filament breakage. Indeed, the incorporation and loss of subunits along pre-existing filaments have been observed in cells (Ngai et al. 1990, Coleman and Lazarides 1992) and shown to occur alongside filament annealing (Çolakoğlu and Brown 2009), but the relationships between these two types of filament dynamics remain unclear. Subunit exchange has been reconstituted *in vitro* using purified vimentin (Nöding et al. 2014). However, the characterization of subunit loss from the lattice was lacking, preventing an accurate estimation of the total number of subunits per filament cross-section and its fluctuations.

The existence of subunit exchange suggests the presence of a pool of soluble subunits in equilibrium with filaments as observed in the cytosol (~1-3 % of total vimentin for cells in culture (Soellner et al. 1985)). However, this pool has been reported as negligible *in vitro* (Sokolova et al. 2006) or undetectable in the absence of phosphorylation (Eriksson *et al*. 2004). In the standard model of IF assembly, tetramers rapidly self-assemble into ULF within seconds when filament elongation is initiated by salt addition, which implies the absence of a significant soluble pool of vimentin (Herrmann and Aebi 2016). Hence, these contradictory observations highlight gaps in our understanding of the vimentin assembly model, and its relationship to subunit exchange. A comprehensive description of vimentin dynamics is needed to elucidate how filament length and the soluble pool of subunits are intrinsically regulated- key factors that determine the network architecture in cells and its potential disruption in disease contexts.

In this study, we elucidate the molecular mechanism responsible for vimentin filament fragmentation by combining *in vitro* single-filament fluorescence and AFM experiments with theoretical modeling. Our findings reveal that filament breakage arises from the constant subunit exchange occurring along the filament length. This subunit exchange leads to local fluctuations in the number of tetramers per cross-section, which weakens the filament and ultimately induces fragmentation. Overall, we demonstrate that the continuous self-repair of filaments through subunit exchange is essential for maintaining their structural integrity, a critical feature for regulating filament length over long time scales.

## RESULTS

### Vimentin subunits spontaneously self-renew along filaments

We propose that filament fragmentation arises from a local, transient dissociation of subunits, which weakens the filament. To test this hypothesis, we first quantified the effect of subunit association/dissociation along the filament over 24 hours-the time scale during which fragmentation occurs under our experimental conditions (Tran et al. 2023). We mixed two populations of filaments: one pre-assembled from recombinant vimentin fluorescently labeled on cysteine-328 and one from unlabeled vimentin (Winheim et al. 2011, Tran et al. 2023, Petitjean et al. 2024)(Fig. 1A). Filaments were imaged at several time points over the 24 hours period, showing that they continued to elongate by end-to-end annealing after mixing as previously described (Tran et al. 2023). End-to-end annealing induced the alternating bright and dim segments observed along the filaments and a continuous increase in filament length (Fig. 1B, Fig. S2).

**FIG. 1:**
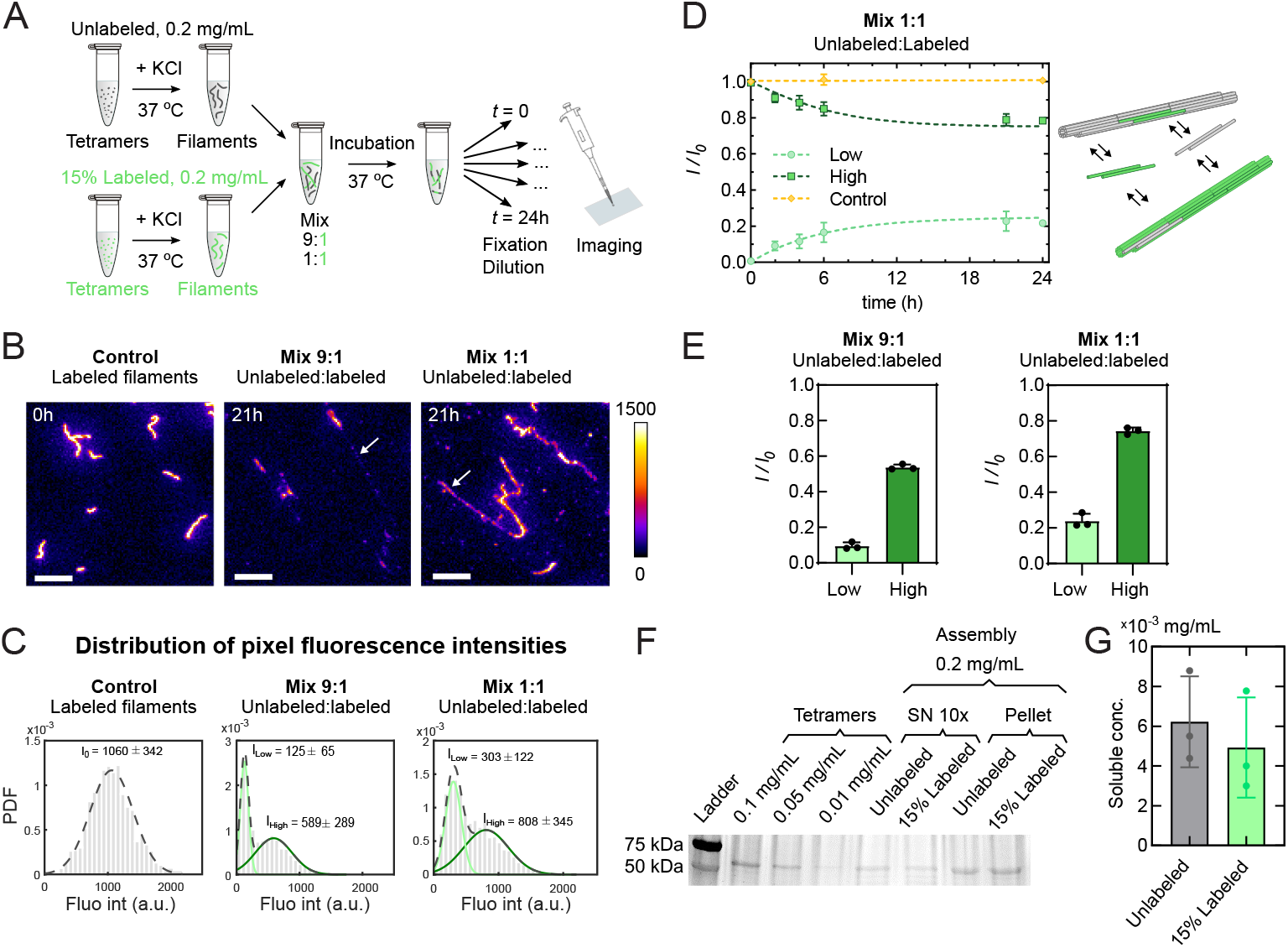
Exchange of subunits between pre-assembled vimentin filaments. (A) Schematics of the subunit exchange experiment between unlabeled and labeled filaments. Two samples of vimentin filaments were pre-assembled by the addition of 100 mM KCl to solutions of tetramers (kick-start method) at an initial concentration of 0.2 mg/mLfor 2h at 37 °C, one without labeling and one with a 15 % AF488 labeling fraction. Then, the two samples were mixed at ratios of 9:1 and 1:1 (unlabeled:labeled) at time 0 and further incubated at 37 °C for up to 24 h. At different time points after mixing, filaments were fixed with glutaraldehyde and diluted 100 times for further imaging. (B) Representative fluorescence images of 15 % labeled filaments at time 0 (control, just before mixing) and filaments 21 hours after mixing at ratios of 9:1 and 1:1 (unlabeled:labeled). The color bar indicates the range of fluorescence intensity. White arrows highlight segments of filaments that were initially unlabeled. Scale bar: 5 μm.(C) Distribution of pixel fluorescence intensities along filaments in the control (15 % labeled filaments at time 0), mix 9:1 and mix 1:1 conditions 21 hours after dilution for one representative replicate. The control distribution was fitted by a single Gaussian to get the mean intensity *I*_0_. The intensity distributions after mixing were fitted with a double Gaussian (dashed line) to determine the mean intensities of filaments initially unlabeled and labeled, referred to as *I*_low_ and *I*_high_ respectively. PDF: Probability distribution functions.(D) Time evolution of the normalized mean fluorescence intensity after filament mixing: initially labeled filaments *I*_high_/*I*_0_ (High, dark green squares), initially unlabeled filaments *I*_low_/*I*_0_ (Low, light green circles) and unmixed labeled filaments *I*(*t*)/*I*_0_ (Control, yellow diamonds). The intensity *I*(*t*) was obtained by a single Gaussian fit of the distribution pixel intensities at time t. Dashed lines: exponential fit with a characteristic rate equal to *k*_off_ = 0.2 *±* 0.1 h^−1^ (SD, *N* = 3) using Eq. 6 (Sup Info), and linear regression (slope = 0.0001 h^−1^) of the control unmixed filaments. Each data point represents the average over *N* = 3 independent replicates (~100 filaments per condition and replicate) and the error bars are the standard deviations of the 3 replicates. Right: Schematics depicting the association/dissociation of vimentin subunits along filament shafts. (E) Bar plot of the normalized intensities Low (*I*_low_*/I*_0_) and High (*I*_high_*/I*_0_) for the ratios of 9:1 and 1:1 21 hours after mixing. Error bars indicate the standard deviation of the 3 replicates.(F) Concentration of soluble vimentin measurement through SDS-page gel: (i) 0.1 mg/mL tetramers; (ii) 0.05 mg/mL tetramers; (iii) 0.01 mg/mL tetramers; (iv) 10× supernatant of unlabeled vimentin, (v) 10× supernatant of 15 % labeled vimentin, (vi) pellet of unlabeled vimentin, (vii) pellet of 15 % labeled vimentin. Vimentin filaments were assembled at 0.2 mg/mL, centrifuged at 140,000 ×*g* for 15 min and the supernatant was concentrated 10 times and compared with a range of known concentrations. (G) Concentration of the vimentin soluble pool, quantified from SDS-page gels. *N* = 3 independent replicates. Error bars are standard deviations.

We evaluated the extent of subunit exchange by quantifying the pixel fluorescence intensities along the filaments. Since the filaments exhibited regions with both bright and dim segments, the pixel intensity distribution for all filaments was bimodal. The first peak corresponded to segments of filaments that were initially unlabeled, while the second peak represented segments that were initially 15 % labeled (Fig. 1C). By fitting the intensity distribution with a double Gaussian function, we estimated the mean intensity of the dim segments (*I*_*low*_) and the bright segments (*I*_*high*_). These values were normalized to the mean fluorescence intensity of the initially 15 % labeled filaments at time 0 just before mixing (*I*_0_, control, Fig. 1C). The normalized mean intensities of the dim and bright segments were then monitored over time during subunit exchange. The time evolution revealed that the mean fluorescence intensity of the dim segments increased over time, concomitant with a decrease in the fluorescence intensity of the bright segments, eventually reaching a steadystate level (Fig. 1D). In control experiments where fluorescent filaments were not mixed with unlabeled filaments, the mean fluorescence intensity remained constant over time (Fig. 1D, control), indicating that the linear density of vimentin subunits per unit length remained stable throughout the assembly process.

If all subunits of the filaments were able to associate/dissociate, we would expect that, in a 1:1 unlabeled:labeled mixture that has reached a steady state, the normalized mean intensity of both the bright and dim segments would reach 0.5, resulting in homogeneous filaments. In a 9:1 unlabeled:labeled mixture, the filaments would also become homogeneous, with a normalized mean intensity of both the bright and dim segments reaching 0.1. However, we observed that the fluorescence distribution remained non-homogeneous along the filaments even after reaching a steady state (Fig. 1B-C, Fig. S2). The normalized mean intensity of the bright segments at steady state reached ~0.75 for a 1:1 unlabeled:labeled mix, and ~0.55 for a 9:1 unlabeled:labeled mix (Fig. 1E). These results indicate that only half of the subunits are exchangeable, while the other half remain immobile. Finally, by modeling the subunit exchange along the filaments using a simple 2-state model (see the theory part in the SI Appendix), we extracted the dissociation rate of the subunit through the exponential fitting of the curves in Fig. 1D, and obtained *k*_off_ = 0.2 *±* 0.1 h^−1^ (mean *±* SD, *N* = 3 independent replicates). We assumed in the model that the subunits were tetramers, which we verified in experiments presented later in this paper. Taken together, our results confirm that there is incorporation and loss of vimentin subunits in already-formed filaments and show that subunit association and dissociation are counterbalanced. Moreover, our results reveal the existence of two populations of subunits: one exchangeable and one immobile, each one accounting for half of the subunits composing the filaments.

### Characterization of vimentin soluble pool

Vimentin filaments spontaneously renew their constituents by association/dissociation of subunits along their length, indicating the existence of a soluble pool of vimentin in equilibrium with the assembled filaments. However, the amount of soluble vimentin is so low that it was barely detectable at physiological ionic strength (100 mM KCl). To detect the soluble pool of vimentin, we first separated polymerized and unpolymerized vimentin by centrifugation at 140,000 ×*g* for 15 minutes, followed by concentration of the supernatant by a factor of 10 using centrifugal filter units. The concentration of soluble vimentin in the supernatant was then measured by quantifying the band intensity on SDS-PAGE gels (SI Appendix). Starting from filaments assembled at 0.2 mg/mL (2 hours at 37 °C), we measured a soluble pool of (5 *±* 2.5) · 10^−3^ mg/mL, which corresponds to 2 % of the total vimentin (Fig. 1F-G). We also compared the concentration of soluble vimentin in both labeled and unlabeled filaments and found no significant difference, suggesting that fluorescent labeling of vimentin has minimal impact on their association/dissociation properties (Fig. 1F-G). Next, we investigated whether a solution of tetramers at the concentration of the soluble vimentin could form filaments by itself using super-resolution microscopy. We performed STochastic Optical Reconstruction Microscopy (STORM) imaging of a solution of 5 × 10^−3^ mg/mL vimentin, 20 % labeled with AF-647, assembled for 6 h at 37 °C and then non-specifically bound to a clean coverslip (Fig. S3A). We did not observe any filaments or filament precursor ULFs, in contrast with control conditions where vimentin filaments were either fully assembled (0.2 mg/mL assembled for 1 h, 40× diluted) or assembled as ULF precursors (0.2 mg/mL assembled for 2 s, 40× diluted) (Fig. S3B). Overall, our results demonstrate that there is a soluble pool of vimentin in equilibrium with filaments. However, the concentration of this soluble pool (~5 × 10^−3^ mg/mL for filaments assembled at 0.2 mg/mL) is so low that it does not form filaments on its own even after 6 hours in the presence of salt.

### Dilution induces both filament thinning and fragmentation

Association/dissociation of vimentin subunits along filament length is taking place in parallel with filament end-to-end annealing/fragmentation. Since dilution of preassembled filaments shifts the balance between elongation and fragmentation towards filament shortening by limiting annealing (Tran et al. 2023), we sought to determine whether dilution would also favor dissociation by decreasing the soluble vimentin pool. To test this, we diluted pre-assembled fluorescently labeled filaments (0.2 mg/mL, 2 h at 37 °C) at different dilution ratios and incubated them further at 37 °C for up to 6 h (Fig. 2A). We imaged the filaments at different time points after dilution and used the mean fluorescence intensity per filament as a measure of the number of subunits per cross-section (Fig. 2B-C). The mean fluorescence intensity and length of the filaments were measured from the fluorescence images using automatic filament detection via the ‘Ridge detection’ plugin in Fiji, with measurements taken along a 2-pixel wide line (Fig.2B). After dilution by 1:200 and 1:500, the mean filament intensity decreased over time, with little difference between dilution ratios, while remaining constant for undiluted filaments (Fig. 2D). After 6 hours of 1:200 dilution, the mean intensity of the filaments reached a saturation value of ~75 % of its initial value, indicating the establishment of a new equilibrium between the soluble pool and the polymerized filaments. At a 1:500 dilution, the filaments became too short after 2 hours for length or intensity to be reliably measured. We verified that the labeling ratio of vimentin did not affect subunit dissociation (Fig. S4), indicating that fluorescent labeling does not impact the subunit exchange process. As previously observed (Tran et al. 2023), the mean filament length decreased with time after dilution due to fragmentation, with a more pronounced decrease at the higher dilution ratio of 1:500 (Fig. 2E).

**FIG. 2:**
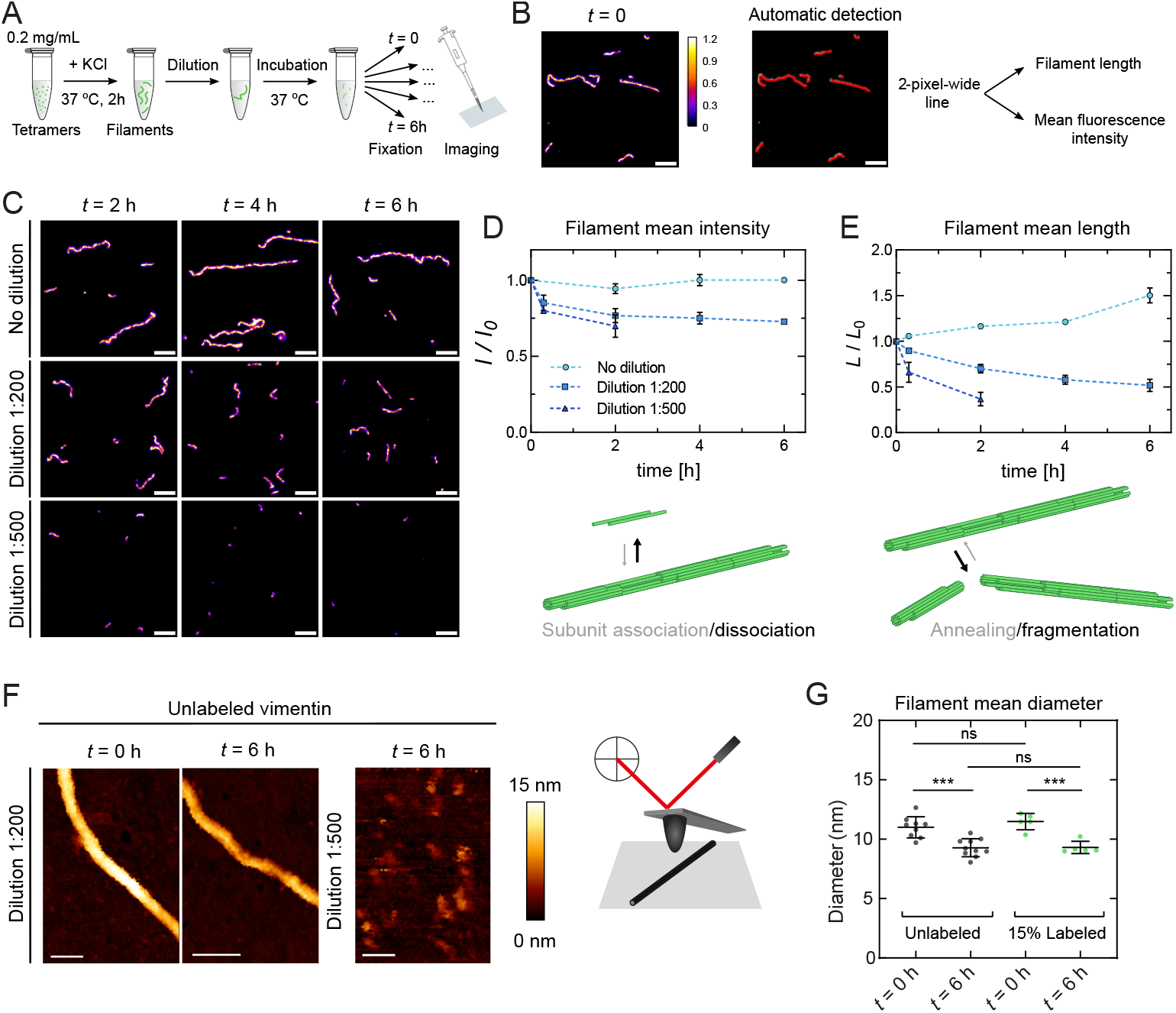
Dilution induces both filament thinning and filament fragmentation. (A) Schematics of the dilution experiments. Vimentin filaments with 15 % AF488 labeling fraction were pre-assembled at an initial concentration of 0.2 mg/mL for 2 h at 37 °C. The assembled filaments were then diluted in the assembly buffer at ratios of 1:200 and 1:500, and further incubated at 37 °C for up to 6 h. Filaments were fixed with 0.25 % glutaraldehyde at different time points after dilution for further microscopy imaging. (B) Representative fluorescence image at time 0, just after dilution 1:200, used as a reference for fluorescence intensity measurements. Scale bar: 5 μm. (C) Representative fluorescence images at time 2h, 4h, and 6h without dilution, after 1:200 and 1:500 dilution. Images were acquired in similar microscopy conditions as in A, and are displayed with the same color scale. Undiluted filaments were diluted 200-fold after fixation to obtain images of similar density to diluted filaments. Scale bar: 5 μm. (D) Time evolution of the normalized mean fluorescence intensity of non-diluted filaments (light blue circles), filaments diluted 1:200 (blue squares) and 1:500 (dark blue triangles). Bottom: cartoon illustrating the unbalanced subunit dissociation associated with the decrease in filament intensity. Filaments diluted 1:500 cannot be detected beyond two hours. (E) Time evolution of the mean length of non-diluted filaments (light blue circles), filaments diluted 1:200 (blue squares) and 1:500 (dark blue triangles). Bottom: cartoon illustrating the unbalanced filament fragmentation associated with decreased filament length. Each data point of (D) and (E) represents the average of *N* = 3 independent replicates (~300 filaments per condition and replicate) and the error bars are standard deviations of 3 replicates. (F) Representative height maps obtained by AFM of unlabeled vimentin filaments immediately after dilution (*t* = 0), 6 hours after dilution at 1:200, and 6 hours after dilution at 1:500. Filaments were fixed with 0.25 % glutaraldehyde and incubated for 5 minutes on a cleaned glass coverslip. The color bar represents the height scale. Scale bar: 50 nm. Right: cartoon of an AFM experiment performed using the Quantitative imaging advanced mode. (G) Scattered dot plot of filament mean diameter for unlabeled vimentin at *t* = 0 and *t* = 6 h after 1:200 dilution (grey dots), and 15 % labeled filaments at *t* = 0 and *t* = 6 h after dilution (green dots). Each dot represents a different filament. *P*-values were calculated with a one-way analysis of variance (ANOVA) performed on the average heights per filament. ***: *P <* 0.001, ns: non significant.

To assess whether the decrease in subunits per cross-section after dilution is linked to filament thinning, we measured filament diameter using atomic force microscopy (AFM). We acquired height maps of unlabeled filaments non-specifically adsorbed on a clean glass coverslip, both immediately after dilution (*t* = 0), and 6 hours after dilution at 1:200 and 1:500 (Fig. 2F). AFM confirmed that no filaments were present after 6 hours at 1:500 dilution. For the 1:200 dilution, the mean filament diameter was calculated by averaging the maximum height values along the filament segment (~300 nm long). We observed a significant 15 % decrease in vimentin mean diameter after 6 hours, also observed for labeled filaments (Fig. 2G). This 15 % decrease in diameter corresponds to a 28 % decrease in cross-sectional area, which aligns with the 25 % decrease in subunits per cross-section measured with fluorescence microscopy. Moreover, the fact that the filament diameter did not depend on whether the filaments were fluorescently labeled or not further demonstrates that the thinning of the filaments after dilution is not a consequence of vimentin fluorescence labeling. Taken together, these results show that filament dilution induces both filament fragmentation and a decrease in subunits per cross-section, leading to filament thinning. This raises the question of whether subunit dissociation could destabilize filament integrity and contribute to fragmentation.

### Addition of soluble tetramers protect diluted filaments from fragmentation

To investigate whether subunit dissociation contributes to filament fragmentation, we conducted experiments in which pre-assembled filaments were diluted into solutions containing vimentin tetramers (Fig. 3A). The aim was to restore the net association of vimentin subunits, counteracting dissociation induced by the low soluble vimentin concentration after dilution and re-establishing the equilibrium between the soluble vimentin pool and polymerized filaments. If filament integrity is preserved under these conditions, susceptibility to fragmentation should decrease. Pre-assembled fluorescently labeled filaments were diluted at ratios of 1:200 and 1:500 into solutions of fluorescently labeled vimentin tetramers at concentrations of 0.1 × 10^−3^, 1 × 10^−3^, and 5 × 10^−3^ mg/mL (Fig. 3A). As previously demonstrated, tetramers at these concentrations do not self-assemble into filaments (Fig. S3). We imaged the filaments at various time points post-dilution and quantified their mean fluorescence intensity and mean length (Fig. 3B-C). Dilution at 1:200 into solutions containing 0.1 × 10^−3^ and 1 × 10^−3^ mg/mL tetramers was insufficient to counteract subunit dissociation and filament fragmentation. In contrast, the presence of 5 × 10^−3^ mg/mL tetramers effectively prevented both effects (Fig. 3B-D). Similar results were obtained with filaments diluted at the ratio 1:500 in 5 × 10^−3^ mg/mL tetramers (Fig. S5). This concentration, corresponding to the soluble vimentin pool (Fig. 1F-G), supports subunit association that balances dissociation from filaments, thereby maintaining constant fluorescence intensity and filament length over time (Fig. 3D, Fig. S5B). To confirm that filament dissociation and rescue were not artifacts caused by fluorescent labeling, we repeated the rescue experiments using unlabeled filaments diluted into solutions of labeled tetramers. Over time, filament fluorescence intensity increased after dilution, confirming that unlabeled tetramers dissociated from filaments and were replaced by labeled ones (Fig. S6). Additionally, when green filaments were diluted into solutions of red tetramers, approximately 25 % of green tetramers were replaced by red ones after six hours of incubation, as indicated by the anticorrelation of red and green signals along the filaments (Fig. S7). These results demonstrate that soluble tetramers maintain filament integrity and that subunit dissociation drives filament fragmentation, proving the tight coupling of the two disassembly mechanisms — filament fragmentation and subunit dissociation.

**FIG. 3:**
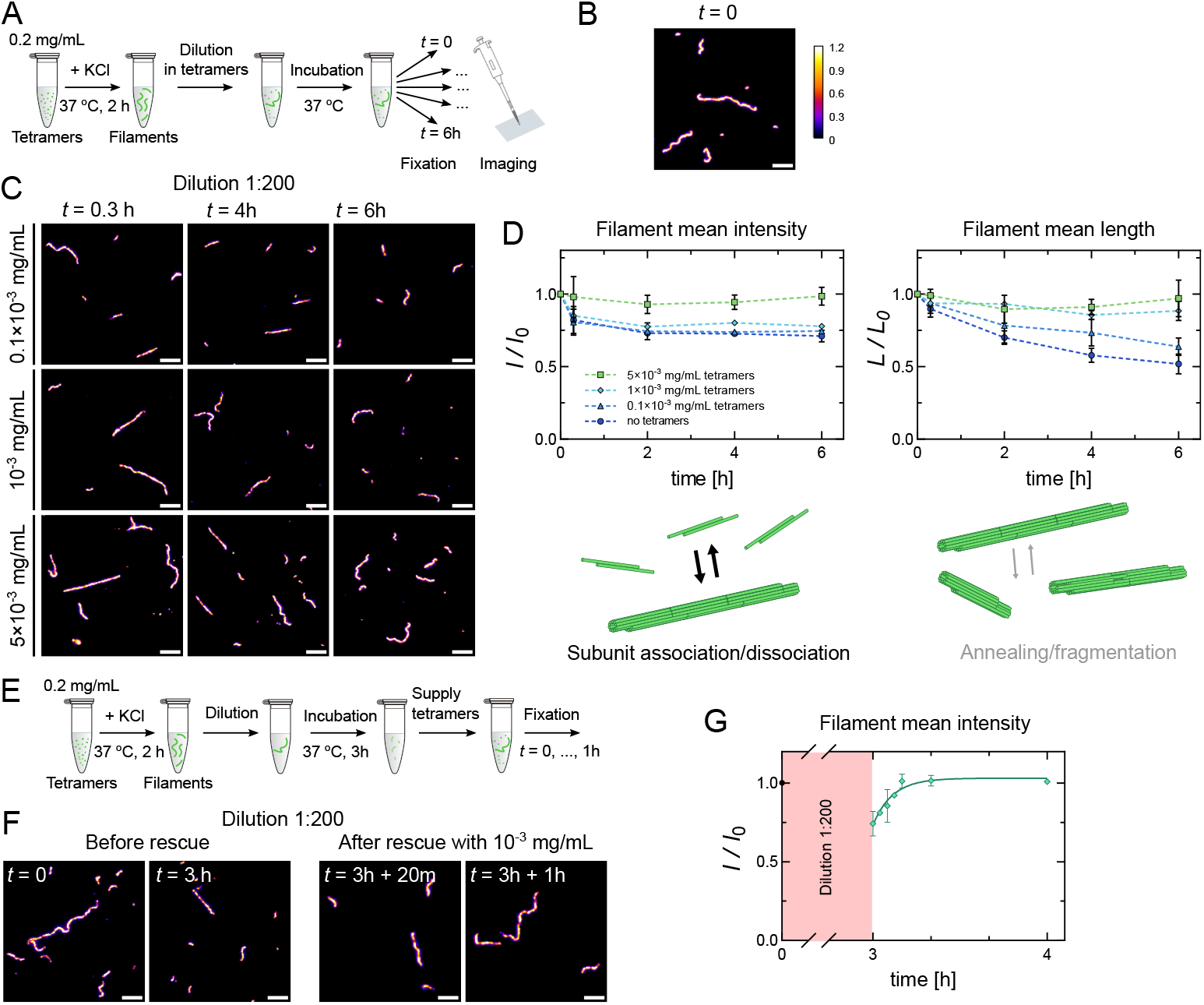
Adding soluble tetramers protects diluted filaments from fragmentation. (A) Schematics of the dilution experiments in tetramers. Pre-assembled vimentin filaments (0.2 mg/mL assembled for 2h at 37 °C) were diluted in a solution containing AF-488-labeled vimentin tetramers at different concentrations in the assembly buffer (0.1 × 10^−3^, 1 × 10^−3^, and 5 × 10^−3^ mg/mL). The mixtures were incubated at 37 °C for up to 6 h. Filaments were fixed with 0.25 % glutaraldehyde at different time points after dilution. (B) Representative fluorescence image at time 0, just after dilution 1:200, used as a reference for fluorescence intensity measurements. Scale bar: 5 μm. (C) Representative fluorescence images at time 0.3h, 4h, and 6h after 1:200 dilution in a 0.1 × 10^−3^, 1 × 10^−3^, and 5 × 10^−3^ mg/mL vimentin solution. Scale bar: 5 μm. (D) Time evolution of the normalized mean fluorescence intensity (left) and mean filament length (right) of filaments diluted 1:200 in a solution without tetramers (dark blue circles), with 0.1 × 10^−3^ mg/mL (blue triangles), 10^−3^ mg/mL (cyan diamonds) and 5 × 10^−3^ mg/mL (green squares). Bottom: cartoon illustrating the subunit dissociation balanced by subunit association when filaments are diluted in a 5 × 10^−3^ mg/mL vimentin tetramer solution, preventing filament thinning. (E) Schematics of filament rescue experiment. Pre-assembled filaments were first diluted 1:200 for 3 h, then supplied with 10^−3^ mg/mL AF-488-labeled tetramers. (F) Representative fluorescence images at time 0h and 3h after dilution before the rescue, and time 20 min and 1 h after the addition of 10^−3^ mg/mL tetramers. Scale bar: 5 μm. (G) Time evolution of the normalized mean fluorescence intensity of filaments to time 0, diluted 1:200 for 3h and then for 2h after the addition of *c* = 10^−3^ mg/mL of tetramers. Solid line: fit by an exponential with a characteristic rate equal to *k*on.*c* up to 1h after the addition of tetramer (see Theoretical modeling in the methods section) Each data point of (D) and (G) represents the average of *N* = 3 independent replicates (~300 filaments per condition and replicate) and the error bars the standard deviation of the 3 replicates.

Next, we examined whether the dilution-induced thinning of filaments was reversible, specifically whether the filaments could regain their initial number of subunits per crosssection. After 3 hours of dilution, tetramers were re-injected into partially disassembled filaments, and their recovery was monitored through imaging at various time points (Fig. 3E-F). When tetramers were added to a final concentration of 10^−3^ mg/mL, the filaments rapidly recovered their initial fluorescence intensity and, by extension, their original number of subunits per cross-section—within minutes (Fig. 3G), demonstrating that filament thinning is reversible. To quantify this recovery process, we applied a 2-state mass conservation model (see theory in the SI Appendix). From the model, the association rate of a subunit into a filament was determined to be *k*_on_ = 9 × 10^3^ (mg.mL^−1^ h)^−1^, equivalent to 0.14 μM^−1^s^−1^ (95 % CI: 0.06–0.28), based on an exponential fit to the data in Fig. 3G. Notably, the equilibrium concentration of the soluble pool is not expected to match the dissociation constant (*k*_off_ /*k*_on_), as this equilibrium depends on the availability of binding sites along the filament shaft, which is influenced by sample preparation and history (see Methods in SI Appendix for details).

### Direct observation of subunit dissociation from the filaments after dilution

Using ensemble measurements and fixing filaments at different time points, we showed that filament dilution induces subunit dissociation, which can be compensated by subunit association if there are enough subunits in the soluble pool. To confirm these results, we designed an *in situ* assay to directly observe subunit dissociation at the level of individual filaments over time. Using Total Internal Reflection Fluorescence (TIRF) microscopy (Fig. 4A), we monitored the changes in the fluorescence intensity of individual AF488-labeled vimentin filaments anchored by antibodies to the surface of a flow chamber. After flowing and incubating the pre-assembled filaments in the chamber, we rinsed with assembly buffer to completely remove unbound filaments and the soluble pool of vimentin. The chamber was sealed and placed on a microscope stage with the temperature maintained at 37 °C. In a standard chamber (height *H* ~100 μm), we observed that the fluorescence intensity of the filaments decreased and reached a plateau at approximately 45 % after 6 hours (Fig. 4B-C), indicating that subunits dissociated from the filaments until a new dynamic equilibrium was reached. This plateau value was below the minimum plateau value of ~75 % observed in bulk experiments (Fig. 2D), suggesting that surface binding of filaments can alter the dynamic properties of their subunits. We verified that photobleaching was negligible as the same number of laser illuminations used to image the control filaments for 6 hours only resulted in less than 5 % intensity loss (Fig. S8). Additionally, the dissociation of subunits from vimentin filaments was unaffected when changing the fluorescence labeling fraction, buffer solutions, or the density of antibodies on the substrate (Fig. S9A-C). This observation indicates that the available surface-anchored antibodies had a negligible effect on the pool of soluble vimentin. The saturation of the fluorescence intensity after 6 hours suggested that the dissociated subunits had formed a new soluble pool in equilibrium with the attached filaments. To test this hypothesis, we performed a second rinse of the chamber to remove all newly dissociated subunits 6 hours after the first rinse (Fig. S10A). The second rinse induced a further decrease in normalized filament intensity (Fig. S10B), confirming that the first plateau was set by the equilibrium between the soluble pool and the filaments. We hypothesized that the fluorescence plateau should be influenced by the chamber volume, which sets the concentration of soluble vimentin and, therefore, the level of filament dissociation. To test this, we repeated the *in situ* dissociation assay using flow chambers of different heights (0.1*H* and 4*H*) (Fig. 4B). Lower-volume chambers resulted in a higher equilibrium intensity plateau, while higher-volume chambers resulted in a lower intensity plateau (Fig. 4C). The relationship between chamber height and plateau value was well predicted by theoretical modeling (Fig. S11), confirming that the chamber volume dictates the concentration of soluble vimentin and the level of filament dissociation. Furthermore, by fitting the fluorescence intensity decay over time with our theoretical description (SI Appendix), we estimated the subunit dissociation rate *k*_off_ = 0.23 *±* 0.08 h^−1^ (mean *±* SD, *N* = 3 independent replicates), which was in good agreement with the value obtained from bulk experiments (*k*_off_ = 0.2 *±* 0.1 h^−1^, Fig. 1). Interestingly, filaments attached to the surface rarely displayed fragmentation after thinning. This may be due to a stabilizing effect induced by the antibodies to which the filaments are attached or because filament breakage is more difficult to detect since the filament tips remain attached to the surface at a distance below the resolution of standard fluorescence microscopy.

**FIG. 4:**
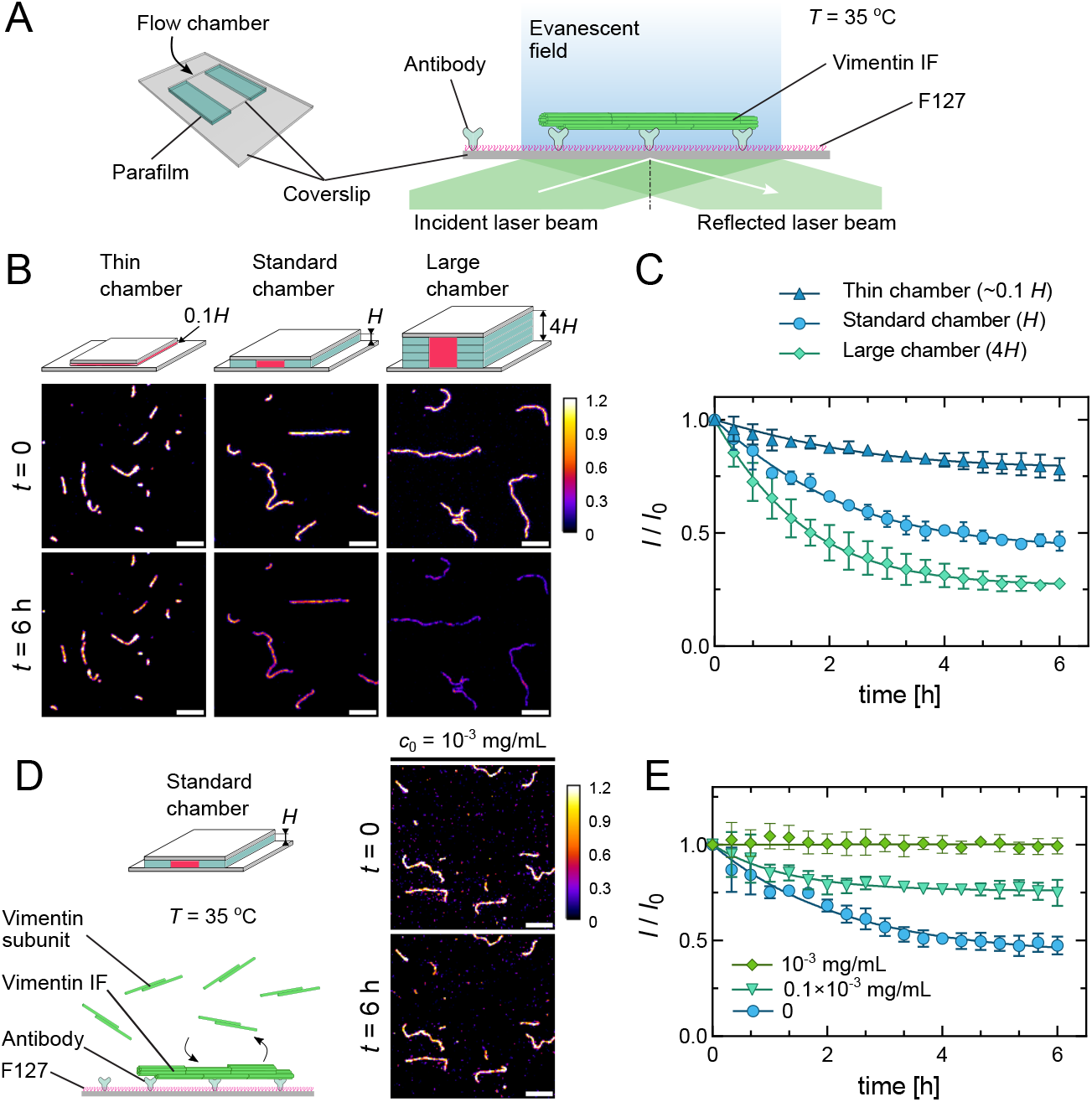
Direct observation of subunit dissociation along filaments after dilution. (A) Schematics of the TIRF microscopy setup for *in situ* filament dilution experiments in a flow chamber. AF-488 15 % labeled pre-assembled vimentin filaments (assembly of 0.2 mg/mL for 2h at 37 °C) were diluted and flushed into a flow chamber made of silanized coverslips, where they can attach to the substrate decorated with anti-AF488 antibodies and passivated by F127. The chamber was rinsed with the assembly buffer to remove all unwanted vimentin subunits, then sealed and incubated at ~35 °C. (B) Schematics of the flow chambers with 3 different chamber heights (0.1*H, H* and 4*H*, with *H* being the thickness of the standard chamber assembled with one layer of parafilm), and representative fluorescence images of filaments at the same observation region, at time 0 vs. 6 h. Scale bar: 5 μm (C) Time evolution of the mean fluorescence intensity of filaments normalized by the value at time 0 and for different chamber heights: 0.1*H* (dark blue triangle), *H* (light blue circles), and 4*H* (green diamonds). Solid line: fit using equation 6 of the SI Appendix. (D) Schematics of the TIRF microscopy setup for *in situ* filament dilution in the standard chamber containing different concentrations of AF-488-labeled tetramers (left). Representative fluorescence images of filaments submerged in 10^−3^ mg/mL tetramers at the same observation region, at time 0 vs. 6 h. Scale bar: 5 μm. (E) Time evolution of the mean fluorescence intensity normalized by the value at time 0, after dilution in the assembly buffer without tetramers supplied (blue circles) and after dilution in a solution of 0.1 × 10^−3^ (light green triangles) and 10^−3^ mg/mL (green diamonds) labeled tetramers. Solid line: fit using equation 6 of the SI Appendix. Each data point of (C) and (E) represents the average of *N* = 3 independent replicates (~50 filaments per condition and replicate) and the error bars are the standard deviation of the 3 replicates.

Next, we tested whether we could also prevent filament thinning in the *in situ* assay, as in bulk experiments, by adding new tetramers after flushing out the soluble pool (Fig. 4D). While the addition of 0.1 × 10^−3^ mg/mL tetramers slowed down global subunit dissociation without completely stopping it, a solution of 10^−3^ mg/mL was sufficient to maintain filament integrity, as filament intensity remained constant throughout the experiment (Fig. 4E). Surprisingly, this tetramer concentration is below the concentration needed to prevent filament fragmentation after dilution in bulk experiments (Fig. 3D, Fig. S5B), suggesting that filament binding to the surface modifies the properties of the filament. In contrast, reinjecting tetramers after thinning only partially restored filament integrity (Fig. S12), unlike when the filaments are free to move in solution (Fig. 3G). The attachment of vimentin filaments on a substrate allowed direct observation of the self-repair mechanism at the single-filament level and confirmed the bulk experiments. It also showed that the attachment of vimentin filaments to the surface affects the overall filament turnover compared to freely diffusive filaments in solution.

### The oligomeric state of the exchanged vimentin subunits is the tetramer

Finally, we aimed to determine the oligomeric state of the subunits in the soluble pool. Previous studies have shown that vimentin phosphorylation induces filament disassembly, with the exchanged subunits being tetramers (Eriksson et al. 2004). Here, we sought to investigate whether the subunits in equilibrium with non-phosphorylated filaments are also tetramers using single-molecule experiments. We used the same standard flow chamber as in TIRF experiments shown in Fig. 4, immobilized labeled vimentin filaments on a substrate, rinsed out the unbound filaments and subunits, sealed the chamber, and incubated it for 60 minutes to allow partial dissociation of subunits from the filaments. Using a microscope with single-molecule fluorescence sensitivity, and a high-power laser illumination setting, we were able to observe multiple fluorescent vimentin subunits diffusing within the chamber. In particular, we focused on the subunits that attached to the substrate and underwent photobleaching (Fig. 5A-B). Fluorescence quantification showed that these subunits (with 15 % labeling fraction) predominantly exhibited a single bleaching step with at most two steps, consistent with the theoretical prediction for tetramers with a 15% labeling fraction (1 fluorophore: 77%, 2 fluorophores: 20%, see details in SI Appendix). We verified that subunits of known oligomeric states and labeling fractions displayed a distribution of bleaching steps consistent with predictions. Immobilized tetramers with 15 % labeling fraction provided a distribution similar to dissociating subunits (Fig. 5C), whereas tetramers with a 50 % labeling fractions showed a higher number of bleaching steps (Fig. 5D). We also verified that the lifetime of a subunit binding to the substrate is longer than its average bleaching time, which ruled out the possibility of subunits/tetramers unbinding from the substrate during the photo-bleaching experiment (Fig. 5E). These results show that the dissociated subunits are tetramers and that the soluble pool of vimentin in equilibrium with the filaments consists of tetramers.

**FIG. 5:**
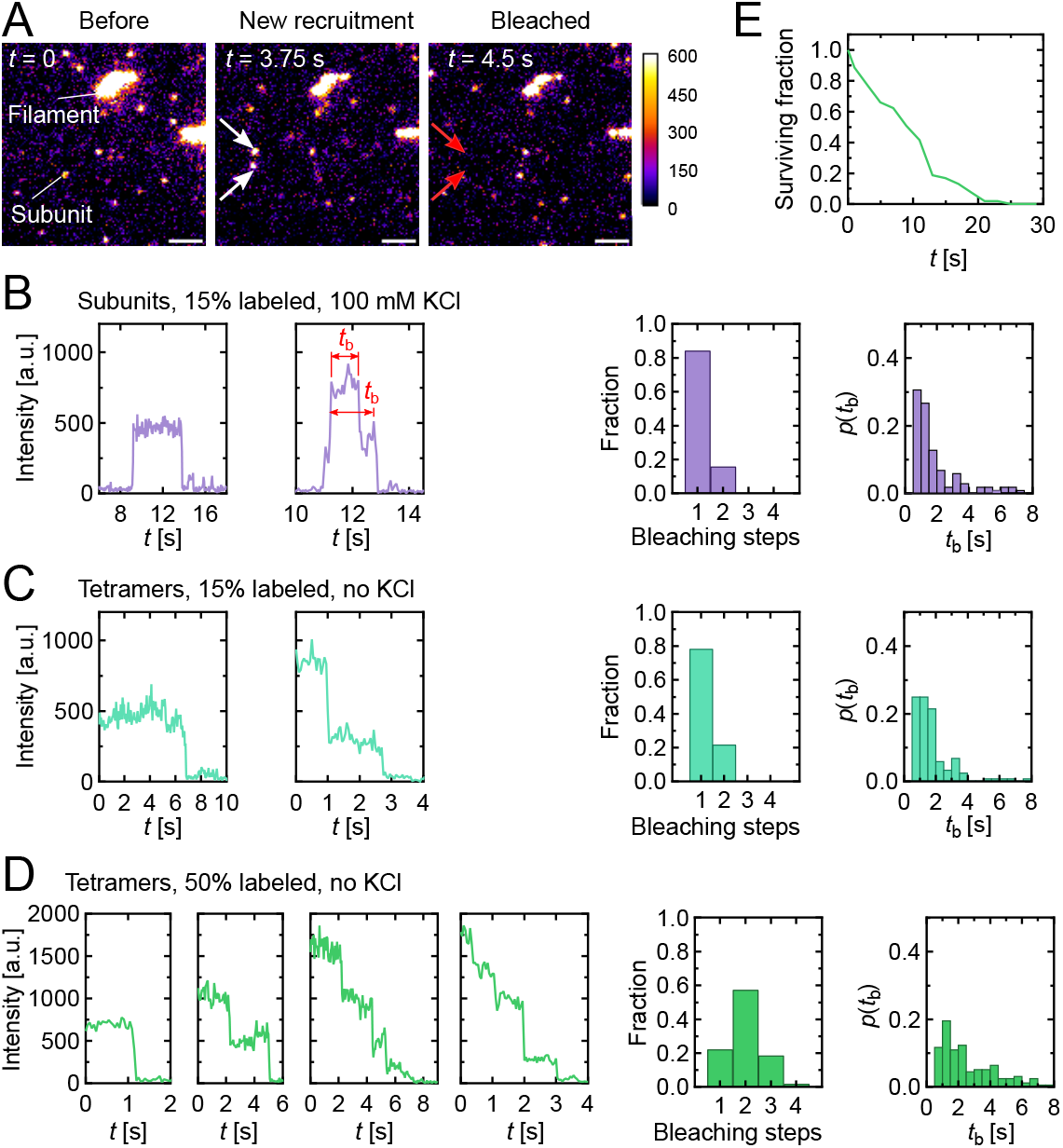
Exchanged subunits are tetramers. (A) Representative fluorescence images of vimentin subunits during bleaching experiments. Pre-assembled filaments (assembly of 0.2 mg/mL for 2h at 37 °C) were introduced to a flow chamber and immobilized on the substrate using the same setup as in Fig. 4A. The flow chamber was rinsed with the assembly buffer to remove all remaining vimentin subunits, then sealed and maintained at ~37 °C. High-power laser was applied to quickly bleach all the subunits dissociating from the filaments and binding to the substrate. White arrows indicate newly recruited subunits on the surface, and red arrows indicate the positions of photobleached subunits. The color scale indicates the fluorescence intensity. Scale bar: 2 μm. The contrast of the images has been set to enhance the visibility of the subunits, resulting in oversaturated-looking filaments. (B-D) Left : examples of fluorescence bleaching steps, middle: distribution of bleaching steps per spot; right: distribution of bleaching times (*t*_b_), in three conditions: (B) 15 % labeled vimentin subunits dissociated from filaments, in assembly buffer; (C) 15 % and (D) 50 % labeled vimentin tetramers, in buffer lacking KCl. Distributions of bleaching steps and durations were obtained from a sample size of ~200 spots for each condition. (E) Fraction of surviving subunits on the surface quantified at low-power laser. The distribution was obtained from 65 subunits.

### Kinetic scenario for filament fragmentation

The quantification of the kinetic parameters pertaining to the association and dissociation of individual subunits allows us to investigate the relationship between these processes and filament fragmentation. We propose that while the detachment of a single subunit does not compromise the integrity of the filament, the simultaneous absence of multiple neighboring subunits can. In a first simple theoretical model, we consider only a cross-section of the filament, which can contain a maximum of 5 mobile tetramers in addition to its immobile tetramers (Fig. 6A). This is consistent with the results presented in Fig. 1, which indicate there is ~50 % of non-exchangeable subunits and with the latest structural data on vimentin showing that there are 10 tetramers per cross-section (Eibauer et al. 2024). We assume that each subunit stochastically associates to and dissociates from the filament independently from the others, as a first approximation, following the rates *k*_on_ and *k*_off_ measured above. As soon as a critical number of subunits *R* are removed from the same cross-section, the filament fragments instantly at this location. By applying first-passage-time theory to a filament in equilibrium with a solution containing a concentration *c* = 100 nM (5×10^−3^ mg/mL) of soluble tetramers (Fig. 1F-G), we compute the average time required for such a fragmentation event to occur, as a function of *R* (Van Kampen 1992) (SI Appendix). We find that the best match to our previous experimental estimate of this time, namely 18 h (Tran *et al*. 2023), is obtained for *R* = 2, and yields a fragmentation time of 63 h.

**FIG. 6:**
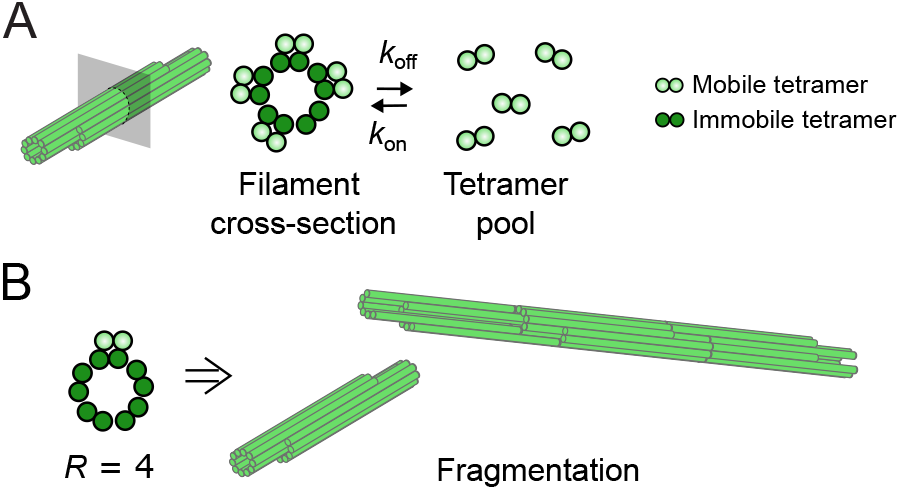
Kinetic scenario for filament fragmentation. (A) Putative model of the two tetramer populations (mobile and immobile) existing in the vimentin filament structure. Immobile tetramers form an inner core, while mobile tetramers form an outer layer and can associate/dissociate with association rate *k*_on_ and dissociation rate *k*_off_. Adjacent tetramers of the outer layer are connected through their tail domains which are not represented in this simplistic cartoon. The exact position of the immobile tetramers within the filament structure is unknown and is hypothetically represented here as an inner layer. (B) Mechanism of filament fragmentation: fluctuations in the number of tetramers per cross-section lead to fragmentation. When the number of mobile tetramers removed reaches the critical value, *R* = 4, the filament breaks at this location.

While this first model yields a fragmentation time in order-of-magnitude agreement with experimental measurements, it also conflicts with our observations in one crucial way. Indeed, the value *R* = 2 implies that a stable filament has at most one mobile subunit missing from each of its cross-sections. This suggests that throughout the disassembly of a collection of fully labeled filaments, no filament with a fluorescence level *I*_high_*/I*_0_ lower than 90 % can be observed. By contrast, in Fig. 3D we observe fluorescence levels as low as 75 %, suggesting that filaments with 2 to 3 missing subunits should still be stable (Fig. 6B). To reconcile our model with this observation, we note that a mobile subunit in a full filament is surrounded on all sides by other mobile subunits and that the resulting binding energy is likely to stabilize it. By contrast, as soon as the filament starts to lose some of its mobile subunits, the remaining subunits tend to lose their stability, which facilitates their departure from the filament. As a coarse model of this facilitation, we propose that the dissociation of the first subunit in a cross-section proceeds with the rate *k*_off_ = 0.2 h^−1^ that we previously estimated (Fig. 1D) for the exchange of subunits between a fully assembled filament and the surrounding solution. Subsequent dissociation events, however, proceed at a faster rate 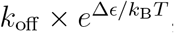, where Δ*ϵ* denotes the difference in binding energy between a fully stabilized and a less-stabilized subunit, and where *k*_B_*T* is the thermal energy. Assuming *R* = 4 compatible with the 75 % intensity level in Fig. 3D, we find that this model achieves a mean fragmentation time of 18 h for a stabilization energy Δ*ϵ* = 4.2 *k*_B_*T*, although choosing *R* = 3 or *R* = 5 each change this estimate by less than 1 *k*_B_*T*. This value of Δ*ϵ* can be interpreted as an estimate of the difference in binding energy between a tetramer that is incorporated into a fully assembled filament, and the binding energy for the same tetramer in a configuration where some of the neighboring tetramers have already been removed.

## DISCUSSION

Here, we identify the molecular mechanism responsible for filament fragmentation using a combination of fluorescence imaging, AFM, and theoretical modeling. Although our fluorescence microscopy assay can only capture the average behavior of tetramer arrival and departure due to diffraction-limited resolution, we were able to quantitatively determine the on-rate and off-rate of individual tetramers along the filaments. First, we establish the existence of a soluble pool of vimentin that is in equilibrium with the assembled filaments. Depleting this soluble pool leads to both filament thinning and fragmentation, whereas the supply of tetramers allows the previously thinned filaments to recover their original state. These results indicate that the soluble pool plays a critical role in filament maintenance by enabling continuous repair, thereby linking the processes of subunit exchange with filament fragmentation.

Our results suggest that the detachment of multiple neighboring subunits is required for fragmentation. We thus develop a theoretical approach that directly exploits dynamical observations and yields insights that are distinct from and complementary to those accessible to structural studies. Our approach robustly rules out a fragmentation mechanism wherein the mobile subunits would be decoupled from one another. We are confident that this conclusion would not change significantly even if we refine our admittedly coarse model of the filament fragmentation geometry. Indeed, any such change would only affect the combinatorial prefactors involved in our transition rates by factors of order one, which would fail to allow for stable filaments with three missing mobile subunits, as observed in Fig. 3. Our second, more sophisticated, theoretical model introduces the possibility that the second, third, etc. subunits are easier to remove than the first. In addition, this model indicates that the binding energy of each of these subsequent subunits is smaller than that of the first by about 4 *k*_B_*T*, consistent with the idea that this binding is mediated by electrostatic interactions or hydrogen bonds. We may further estimate the binding energies of the subunits by interpreting their dissociation time 1*/k*_off_ as the time required for a diffusing rod of diameter *d* = 2 nm and length *l* = 50 nm to jump over an energy barrier whose height *ϵ* equals the binding energy of the subunits with the rest of the vimentin network. This simple model implies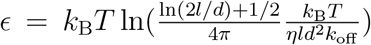, where *η* = 10^−3^ Pa.s 4*π ηld*2*k*_off_ is the viscosity of water. Numerically, this expression yields a binding energy of a mobile subunit fully surrounded by other mobile subunits of the order of *ϵ* ≃ 26 *k*_B_*T*. This energy is reduced by an amount of order Δ*ϵ* = 4 *k*_B_*T* when one of these surrounding subunits is removed. While our model makes the simplifying assumption that the removed subunit is in the same filament cross-section as the subunit of interest, similar conclusions could be drawn from a model considering the removal of neighboring subunits along the filament’s longitudinal axis. Our value of Δ*ϵ* should thus be understood as a rough first estimate to be refined as the geometry of the fragmentation problem becomes better understood.

We have identified two distinct populations of tetramers within the filament: exchangeable and immobile ones. Based on the cylindrical geometry of the filaments, we hypothesize that the immobile tetramers are localized at the center of the cylinder, forming an inner core, whereas the exchangeable tetramers are positioned on the outer layer, facilitating their exchange with the bulk (Fig. 6A). This model of tetramer organization is intuitive but remains speculative, as our fluorescence measurements currently lack the resolution to precisely determine the localization of tetramers within the filament cross-section. The existence of two distinct populations of tetramers also does not align with the atomic model of vimentin derived from cryoEM imaging and recently published by Eibauer *et al*. (Eibauer et al. 2024), in which all tetramers are equivalent. In the Eibauer model, each tetramer contributes to both the inner and outer layers depending on the specific cross-section along the filament.

However, since amino acid side chains are not visible in the 7.2-*Å* resolution cryoEM density maps, the contribution of individual residues to the mechanism of vimentin assembly cannot be discerned at this time (Lomakin et al. 2024). Although our results appear inconsistent with the atomic model, they may still align with the density map. Further investigations, including cryoEM imaging of diluted filaments, are needed to gain a deeper understanding of the detailed organization underlying the distinct dynamic properties of these two tetramer populations.

Previous bulk experiments have shown that the way vimentin filaments are assembled (kick-start *vs*. dialysis) impacts subunit exchange (Nöding et al. 2014). “Kick-start” refers to the triggering of filament assembly by the addition of 100 mM KCl in one go, while “dialysis” refers to the gradual increase of the KCl concentration from 0 to 100 mM during the dialysis process. Slow assembly has been shown to decrease the variability in the number of subunits per cross-section along filaments, a phenomenon known as polymorphism (Herrmann *et al*. 1996). We quantified the impact of the assembly method on subunit dissociation in the *in situ* assay and observed that filaments, which are less polymorphic, exhibit slower dissociation (Fig. S9D). This is consistent with the fact that tetramers with fewer neighboring subunits are more prone to detachment. Additional experiments would be needed to further characterize the difference in binding energy between polymorphic filaments and those assembled by dialysis. For instance, single-molecule experiments could be conducted using optical tweezers to measure the force required to remove a tetramer from the filament shaft, as has been quantified for microtubules (Kuo et al. 2022). The effect of filament thinning after dilution or phosphorylation on this removal force could also be investigated.

The soluble pool of vimentin tetramers in equilibrium with the filaments allows the filaments to maintain their integrity. In a cellular context, there may be unidentified proteins that interact with soluble vimentin subunits and sequester them. Sequestration of subunits would prevent continuous self-repair of the filaments, thereby promoting filament fragmentation, which could serve as a mechanism to regulate filament length. Subunit sequestration has been shown to regulate the dynamics of other types of cytoskeletal filaments (Belmont and Mitchison 1996). In the same line of thought, the recruitment of proteins to the surface of the filaments may also stabilize or destabilize the filaments by modulating the subunit association/dissociation rates. Another way to regulate filament-subunit exchange in cells is by modifying the subunit association/dissociation rates through changes in the binding energy between the subunits. Previous studies have shown that vimentin phosphorylation triggers filament disassembly (Inagaki et al. 1987, Eriksson et al. 2004, Snider and Omary 2014). Eriksson *et al*. showed that vimentin phosphorylation increases the soluble pool of tetramers *in vivo* and *in vitro* (although the *in vitro* soluble pool in the absence of phosphorylation was previously undetectable), suggesting that vimentin phosphorylation changes the equilibrium constant of subunit exchange towards a higher off-rate (Eriksson et al. 2004). Further work is required to characterize the effect of phosphorylation, quantify the modified on- and off-rate, and verify whether phosphorylation promotes filament thinning and fragmentation. Other PTMs could also be investigated, such as S-gluathionylation, which has been shown to induce vimentin fragmentation (Kaus-Drobek et al. 2020), or oxidation, which induces vimentin remodeling in biomolecular condensates (Martínez-Cenalmor *et al*. 2024). Finally, another way to tune the IF length and network topology through variation in subunit association/dissociation would be to change the filament composition. IFs represent a large family of proteins (*>* 70 genes in humans). Depending on the type of cell and tissue, as well as differentiation and stress, different IF proteins can co-assemble within the same filament, leading to a complex IF composition (Leduc and Etienne-Manneville 2015). Subunit exchange could be a way to modify filament composition progressively without completely disassembling the IF network.

Vimentin filaments, whether freely diffusing in solution or bound to a surface, show little difference in the rate of dissociation of tetramers from the filament. However, tethering the filaments to the surface does affect the overall turnover. Attached filaments exhibit stronger thinning after the removal of soluble vimentin and require a 5-fold lower concentration of tetramers in solution to maintain their integrity compared to filaments in solution. Furthermore, thinning can only be partially reversible for filaments attached to the surface (Fig. S12). We propose that this difference between bulk and *in situ* experiments may be partly due to stabilization effects induced by anchoring filaments to the surface via antibodies. Binding to the surface may preserve the integrity of the filaments by preventing fragments from drifting apart. Furthermore, anchoring the filament could induce internal reorganization of the filament, as previous studies have shown that vimentin filaments flatten when attached to a surface (Lorenz et al. 2019).

Experiments on purified vimentin filaments have shown that individual filaments can be stretched up to 350 % without breakage (Block et al. 2017). These remarkable mechanical properties are attributed to the hierarchical organization of the tetramers within the filaments. For strain up to ~100 %, vimentin stretching results from the unfolding of its alpha-helices (Nunes Vicente et al. 2022, Lorenz et al. 2019, Block et al. 2018, Forsting et al. 2019). Interestingly, vimentin filaments keep softening upon repeated cycles of stretching and relaxation, indicating that filament mechanics depends on the strain history. It would be interesting to probe the impact of filament stretching on the association/dissociation rate of tetramers. One could also test whether subunit exchange could reset filament mechanics by providing new tetramers to stretched filaments and allowing filament renewal through incubation at 37 °C for a few hours. We believe that conducting experiments at 37 °C is fundamental for a complete understanding of the relationship between mechanical and dynamic properties. The partial reset of filament could also be relevant in cells, where the soluble pool of vimentin can be regulated. Thus, the self-repair mechanism uncovered in this work may have broader implications for cellular functions involving vimentin, such as cell resilience, resistance to large deformations (Hu et al. 2019), or mechanosensitivity (Ndiaye et al. 2022). Interestingly, the soluble pool of cellular vimentin has been shown to be sensitive to stimuli that alter cellular tension and morphology (Murray et al. 2014).

Filament self-repair has also been observed in another type of cytoskeletal filament, namely microtubules. Recent studies have shown that tubulin dimers can be exchanged along the filament shaft, and this self-repair mechanism promotes filament rescue (Théry and Blanchoin 2021). The renewal of tubulin dimers is spontaneous and occurs at lattice defects or damage sites, impacting both the dynamical and mechanical properties of microtubules (Théry and Blanchoin 2021). In the absence of prior defects, estimates of the free energy required for a tubulin dimer to dissociate from the (intact) microtubule lattice range from 35 to 80 *k*_B_*T* (Schaedel et al. 2019), suggesting that spontaneous exchange of tubulin is negligible over the lifetime of the microtubule (a few minutes). In contrast, since the lifetime of vimentin filaments is much longer (a few tens of hours) and the free energy of vimentin tetramers is much smaller (~30 *k*_B_*T*, as measured here), spontaneous vimentin subunit exchange plays a critical role. We have shown here that the self-repair mechanism is important for regulating filament dynamics, but the link between self-repair and mechanical properties remains to be established.

In conclusion, we reveal a novel self-repair mechanism, in which the exchange of tetrameric subunits maintains vimentin filament integrity. These findings unveil the essential role of the soluble pool. Importantly, we also identify two distinct populations of tetramers within the filaments, raising new questions regarding the intermediate filament architecture.

## METHODS

Detailed protocols are included in the SI Appendix. We purified recombinant wild-type human vimentin from bacteria and labeled it using Alexa Fluor (AF) 488, AF 568, and AF 647 maleimide dyes that form covalent bonds with the cysteine-328 on vimentin as described previously (Winheim et al. 2011, Tran et al. 2023). Vimentin, stored in 8M urea at −80 °C, was renatured by stepwise dialysis to sodium phosphate buffer (pH 7.0, 2.5 mM sodium phosphate, 1 mM DTT). We typically assembled vimentin filaments using a standard ‘kick-start’ assembly protocol by adding 100 mM KCl (final) to a solution of vimentin at the desired concentration and labeling rate, typically 0.2 mg/mL with 15 % labeling, and incubating the mix at 37 °C for 2 h. For the bulk experiments, filaments were fixed at different time points using 0.25% glutaraldehyde, diluted 100 times for non-diluted samples, and observed using standard epi-fluorescence microscopy. To observe filaments thinning *in situ*, we followed a protocol of TIRF microscopy experiment from our previous works (Tran et al. 2023, Petitjean et al. 2024) where filaments are attached to the coverslips of a flow chamber, functionalized with antibodies anti-AF488.

## Supporting information

Vimentin_Revision_bioRxiv.pdf

## ACKNOWLEDGMENT

We thank the Romet-Lemonne/Jegou lab for assistance and exciting discussions, and Ingrid Billault-Chaumartin for careful reading of the manuscript. CL thanks Harald Herrmann and Tatjana Wedig for teaching her vimentin purification, and the support of the EU-supported Cooperation in Science and Technology (COST) action NANONET. This project was funded by “Investissements d’Avenir” Labex WhoAmI? (ANR-11-LABX-0071) and the Université Paris Cité IdEx (ANR-IDEX-0001), LabEx PALM (ANR-10-LABX-0039-PALM), the Agence Nationale pour la recherche (ANR-21-CE11-0004-02), the Impulscience program of Fondation Bettencourt-Schueller and the Centre National de la Recherche Scientifique.

## Notes

### Competing Interest Statement

The authors have declared no competing interest.

### Summary of Updates

We have added new AFM data to Fig 2, which provides new experimental evidence of filament thinning after dilution independent from fluorescence measurements. We have also rewritten the discussion part.

