## Supplementary material for "Continuous self-repair protects vimentin intermediate filaments from fragmentation": Vimentin_Revision_bioRxiv.pdf

#### This PDF file includes:

Supporting text
Figs. S1 to S12
SI References

### Supporting Figures

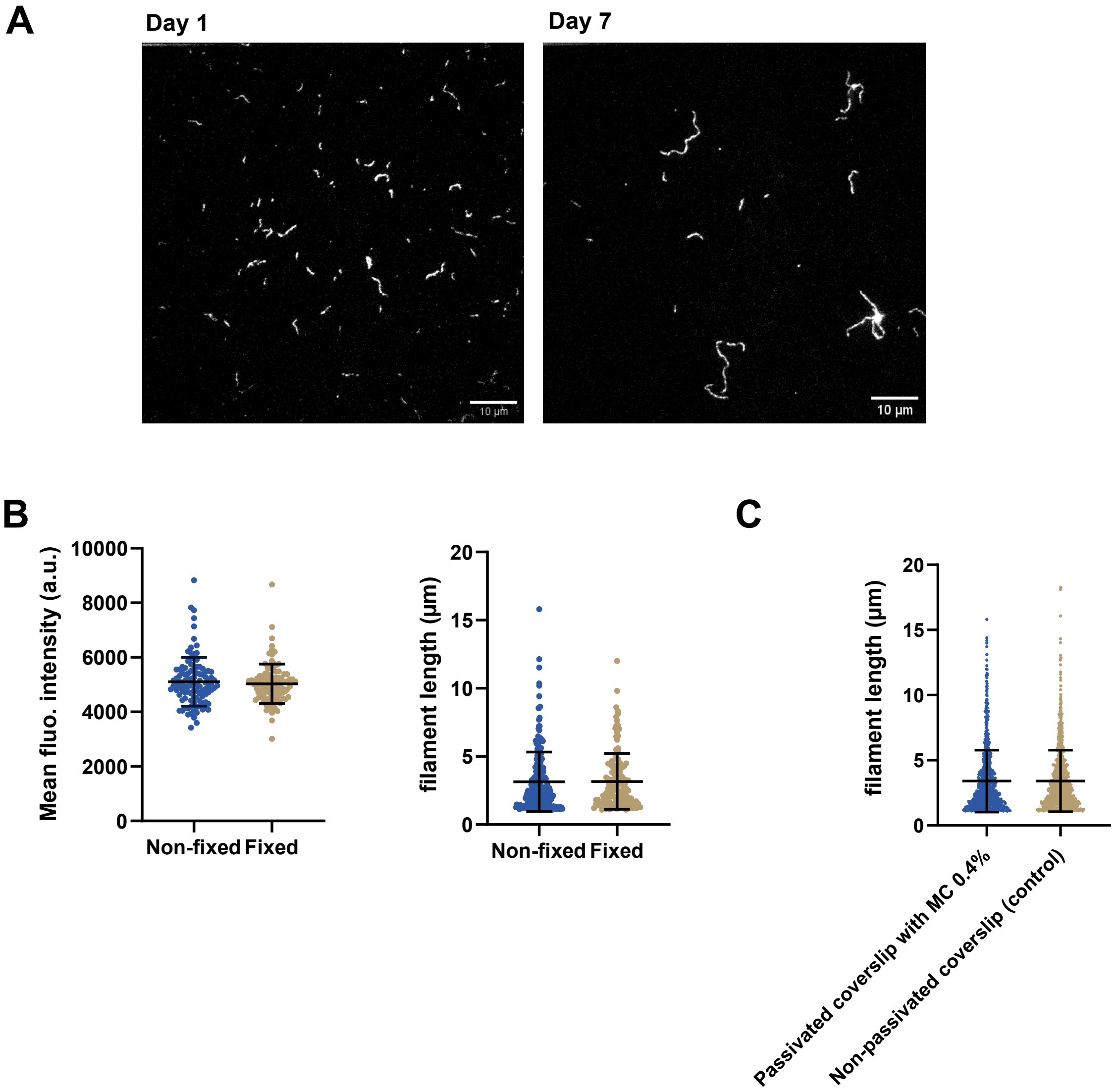


**Fig. S1.** Controls for vimentin imaging. (A) Representative fluorescence images of vimentin filaments grown for 2 hours at 37 ℃ on day 1 (left) and after 7 days spent on the bench (right). (B) Comparison of the mean fluorescence intensity (left) and filament length (right) without (blue dots) and with fixation with 0.25% glutaraldehyde (brown dots). About 100 filaments were analysed, and no significant difference was observed. Filaments grown for 2 hours at 37 ℃ were diluted 100 times and observed immediately after dilution for the "non-fixed" condition; or fixed with 0.25% glutaraldehyde, diluted 100 times and imaged for the "fixed" condition. (C) Distribution of filament length when observed in a thin chamber made of non-passivated glass coverslips (control, standard procedure used for Fig. 1 to 3) (brown dots) and in a flow chamber made of coverslips passivated by DDS and F127 (as in Fig. 4)(blue dots). Filaments were fixed and diluted 100 times for standard observation in a thin chamber, and fixed, diluted 100 times, and supplemented with 0.4% methylcellulose for observation in the passivated flow chamber. Methylcellulose allowed the filaments to be forced close to the glass surface for observation without being attached to the glass. No significant difference was observed between the two distributions indicating that filament attachment to the glass coverslips for observation has minimal impact on the estimation of the mean filament length. Error bars in (B-C) indicate the mean and standard deviation.

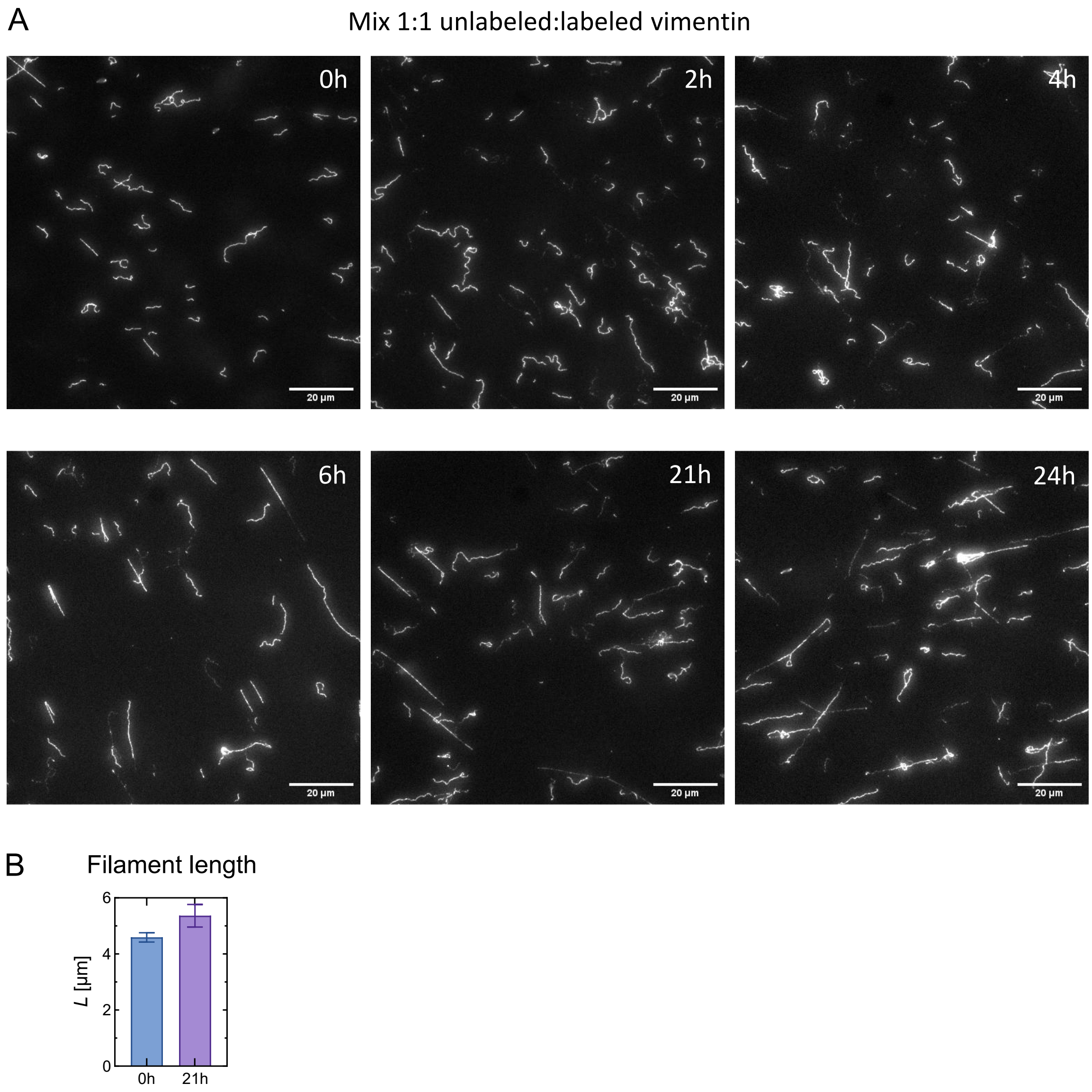


**Fig. S2.** Time evolution of subunit exchange after mixing of preassembled unlabeled and 15% labeled filaments at a ratio 1:1. (A) Representative fluorescence images of mixed filaments at different time points after mixing. The time after mixing is written on the top right corner of the images. (B) Mean length of the mixed filaments at 0 h and 21 h after mixing. Error bars are standard errors of the means over 200 filaments.

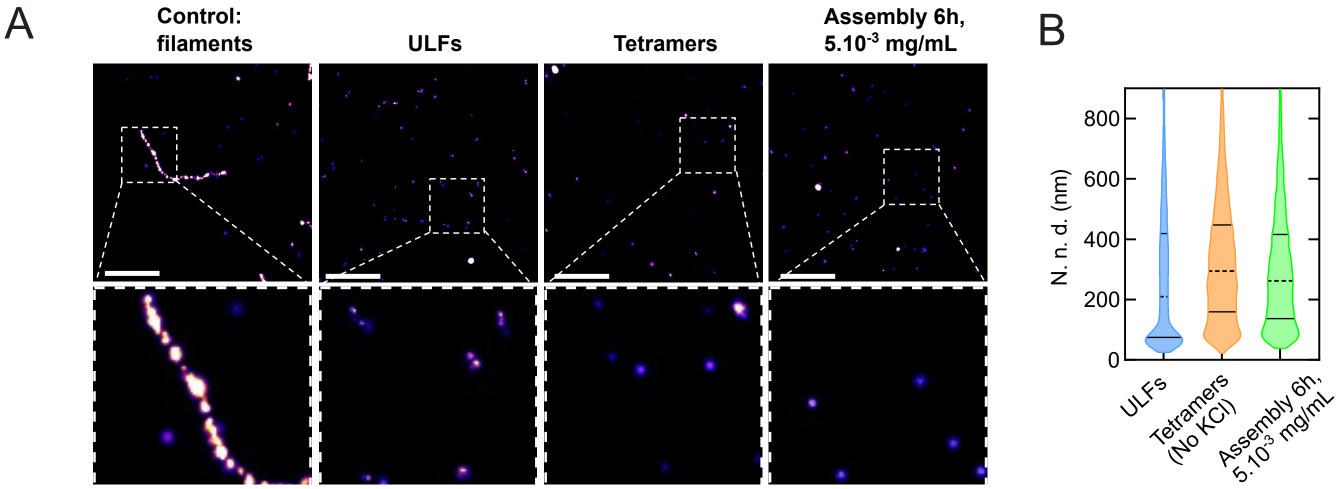


**Fig. S3.** Vimentin at $5 \times 10^{-3}$ mg/mL cannot form filaments. (A) Representative 2D STORM images of vimentin filaments (assembly at 0.2 mg/mL for 1 hour with 100 mM KCl, then diluted 20x), ULFs (assembly at 0.2 mg/mL for 2 seconds with 100 mM KCl, then diluted 20x), tetramers (no assembly, vimentin at 5.10-3 mg/mL) and vimentin assembled at a concentration of $5 \times 10^{-3}$ mg/mL for 6 hours. Inset: zoom of the boxed region. The four conditions were acquired with 20 % vimentin labeled with AF 647. Scale bar: 1 $\mu$m (top images) and 250 nm (bottom images). (B) Violin plot of the nearest neighbor distances (N. n. d.) for the conditions ULFs, non-assembled tetramers, and $5 \times 10^{-3}$ mg/mL vimentin assembled for 6 hours. Quantification was performed on three images of $40 \times 40$ $\mu$m for each condition, with 9000 to 10 000 n.d.d per condition. The medians are represented by dashed lines, and the quartiles are depicted as solid lines.

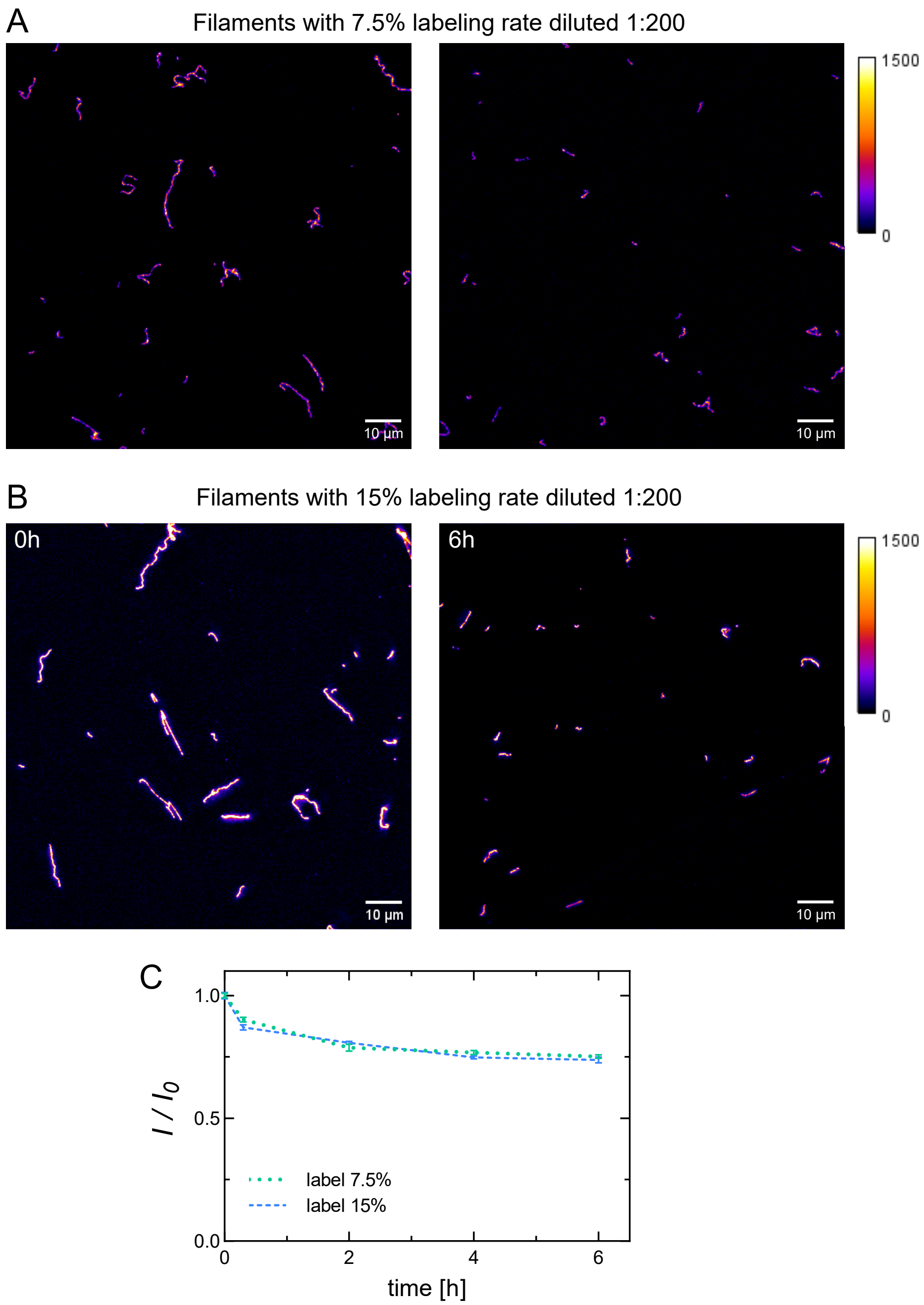


**Fig. S4.** Vimentin fluorescent labeling does not affect subunit dissociation. Comparison between 7.5 % and 15 % labeling rate. (A) Representative fluorescence images of vimentin filaments with 7.5% labeling rate and (B) 15 % labeling rate, diluted 1:200 in assembly buffer and incubated at 37 ℃ for 0 h and 6 h. The color bar indicates the range of fluorescence intensity of the filaments. Scale bar: 10 µm. (C) Time evolution of the mean fluorescence intensity normalized by the intensity value at time 0, after 1:200 dilution in assembly buffer. Each time point is an average of over ∼300 filaments, and the error bars are standard errors over ∼300 filaments.

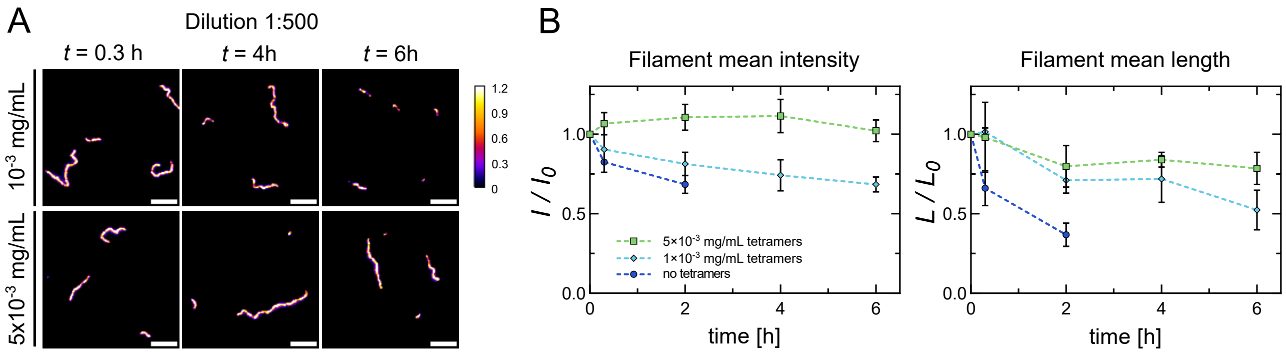


**Fig. S5.** Adding soluble tetramers protects diluted filaments from fragmentation at ratio 1:500. (A) Representative fluorescence images at time 0.3h, 4h, and 6h after 1:500 dilution in a $10^{-3}$ and $5 \times 10^{-3}$ mg/mL vimentin solution. Scale bar, 5 $\mu$m. (B) Time evolution of the normalized mean fluorescence intensity (left) and mean filament length (right) of filaments diluted 1:500 in a solution without tetramers (dark blue circles), with $10^{-3}$ mg/mL (cyan diamonds) and $5 \times 10^{-3}$ mg/mL (green squares). Each data point represents the average of 3 independent replicates (about 300 filaments per condition and replicate) and the error bars the standard deviation of the 3 replicates.

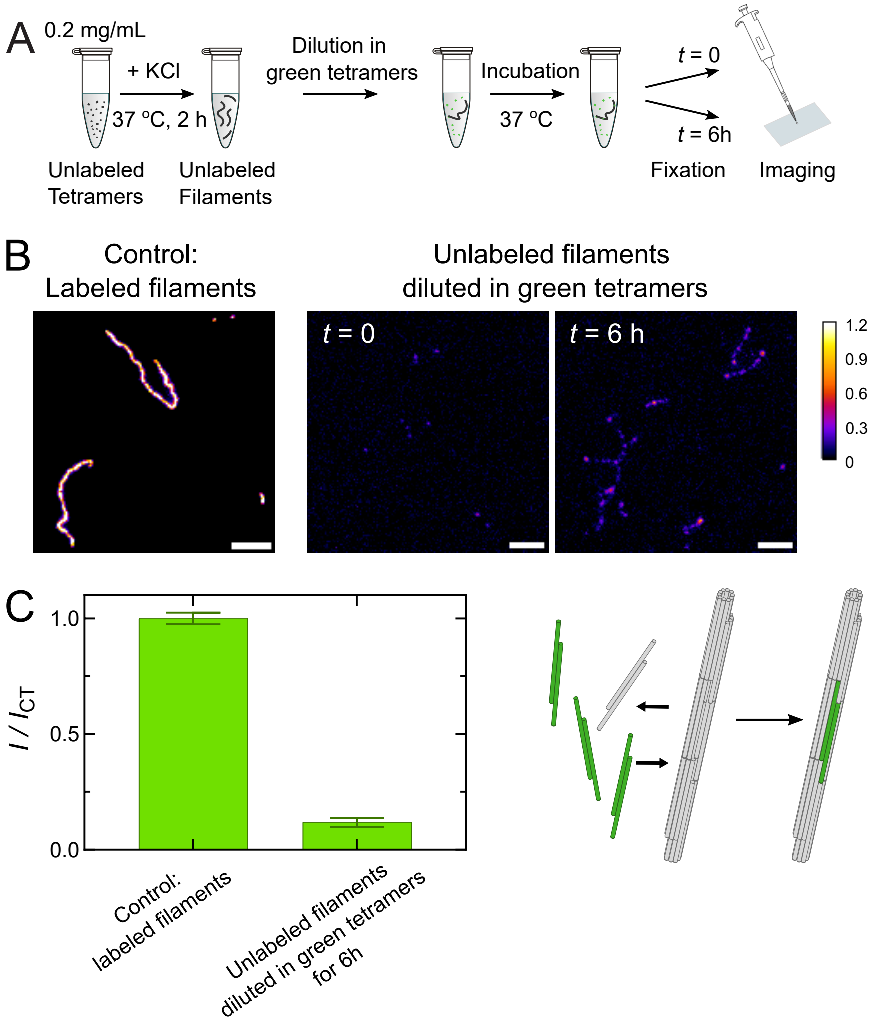


**Fig. S6.** Dilution of unlabeled filaments in $10^{-3}$ mg/mL labeled tetramers. (A) Schematics of the 1:200 dilution experiments of unlabeled filaments in $10^{-3}$ mg/mL green tetramers. (B) Fluorescence images of labeled filaments (control assembly) assembled from the same labeled tetramers used in the dilution experiment (0.2 mg/mL, 15% labeling fraction, assembly for 2 h at 37 °C) (left), and images of unlabeled filaments after diluted in labeled tetramers at time 0 vs. 6 h (right). Scale bar: 5 $\mu$m. (C) Quantification of mean fluorescence intensity along unlabeled filaments at time 6 h, normalized by the average intensity of the labeled filaments in the control assembly (left). Schematic illustrates the outcome of the experiment with labeled subunits associated with unlabeled filaments (right). The error bars are standard errors over 100 filaments.

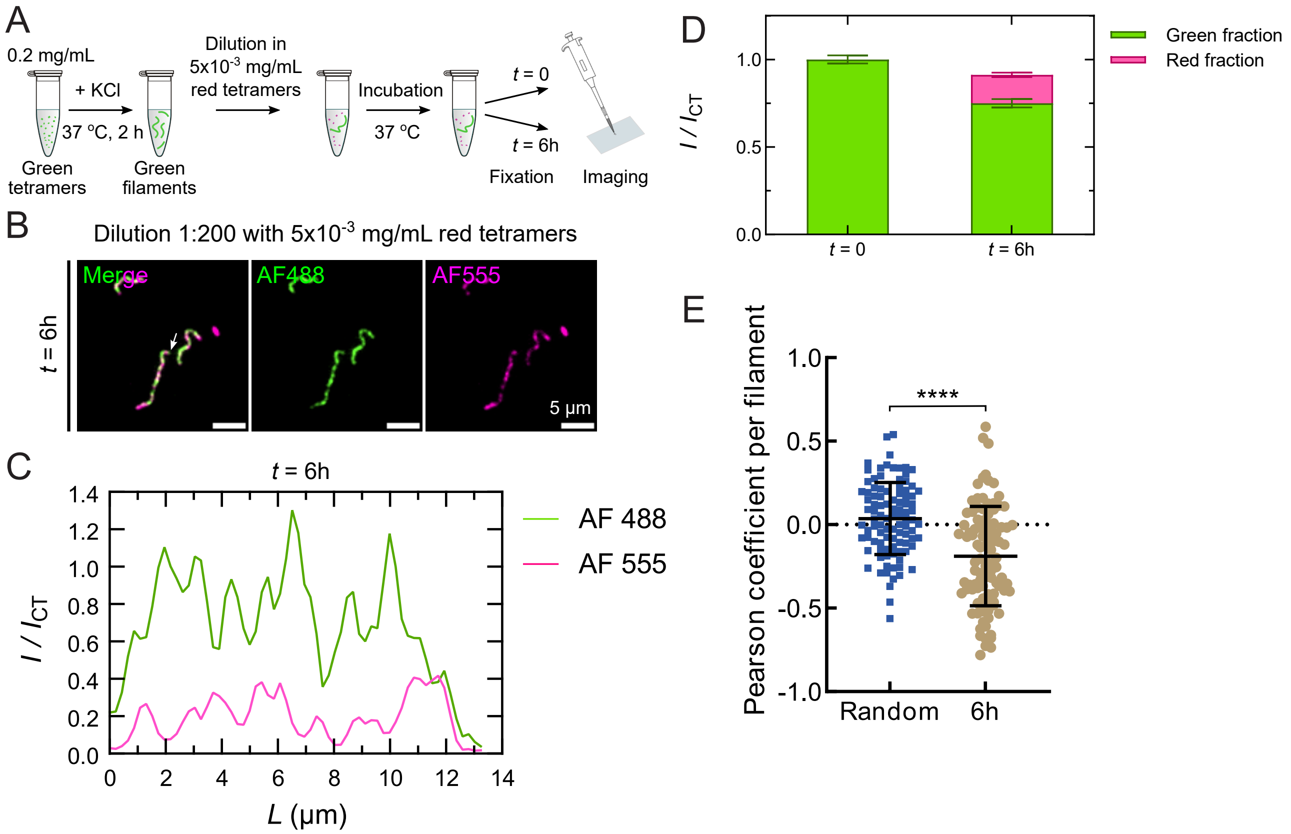


**Fig. S7.** Dilution of green filaments in red tetramers shows that the red tetramers are added at the position where green tetramers have dissociated. (A) Schematics of the 1:200 dilution experiments of green labeled filaments in $5 \times 10^{-3}$ mg/mL red tetramers (B) Fluorescence images at time 0 and 6h in the green and red channels, and overlay. Scale bar: 5 $\mu$m (C) Fluorescence intensity profiles of the filament indicated by the white arrow in (B) at 6h in the green and red channels. The green signal is normalized by the mean intensity value along green filaments before dilution at time 0 ($I_{\mathrm{CT}}$) and the red signal by the mean intensity value along control red filaments grown only with red tetramers for 3h at 0.2 mg/mL ($I_{\mathrm{CT}}$). (D) Quantification of the mean fluorescence intensity along the filaments before (t=0) and after dilution (6h) for $N = 100$ filaments. The mean green intensity is normalized by the mean value at $t = 0$ ($I_{\mathrm{CT}}$). The red intensity is normalized by the mean intensity value along red filaments grown only with red tetramers for 3h at 0.2 mg/mL. The error bars are standard errors over 100 filaments. The green signal at 6h corresponds to 75-80% of the signal at time 0, indicating that 25% of the green subunits have been exchanged with red subunits. Dilution in $5 \times 10^{-3}$ mg/mL tetramers prevents filament thinning. (E) Quantification of the anti-correlation between the green and the red signal along vimentin filaments 6h after 1:200 dilution of green filaments in $5 \times 10^{-3}$ mg/mL red tetramers. Each dot corresponds to the Pearson correlation coefficient for a single filament, comparing the fluorescence intensity of each pixel in the green vs red channels. In control conditions ("Random" on the graph), the position of the red pixels was randomly mixed. We used a paired $t$-test to compare the distribution. ****: $p < 0.0001$,.

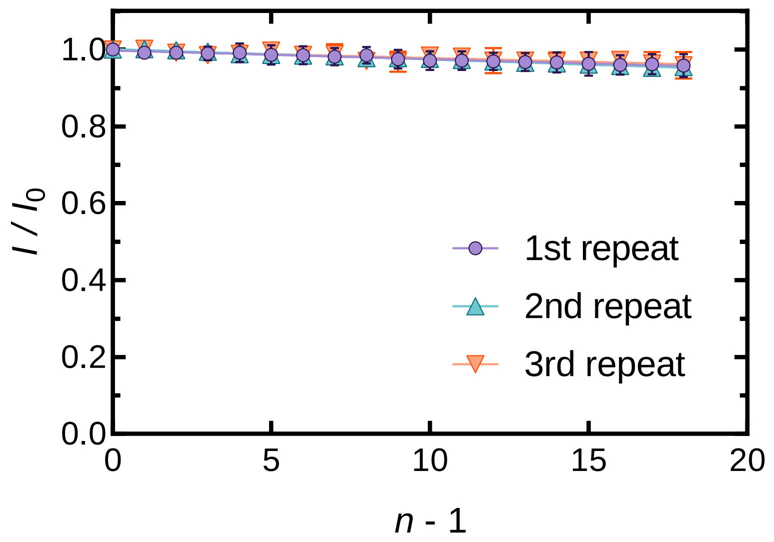


**Fig. S8.** Bleaching control of the labeled filaments on the substrate in TIRF microscopy experiments. Mean fluorescence intensity of vimentin filaments normalized by the intensity value at time 0, over $n$ illuminations, acquired at 1 illumination per 2 s. This bleaching control was performed right before each TIRF experiment. The graph shows 3 representative replicates, where each time point represents an average over $\sim$50 filaments and the error bars are standard deviations over $\sim$50 filaments.

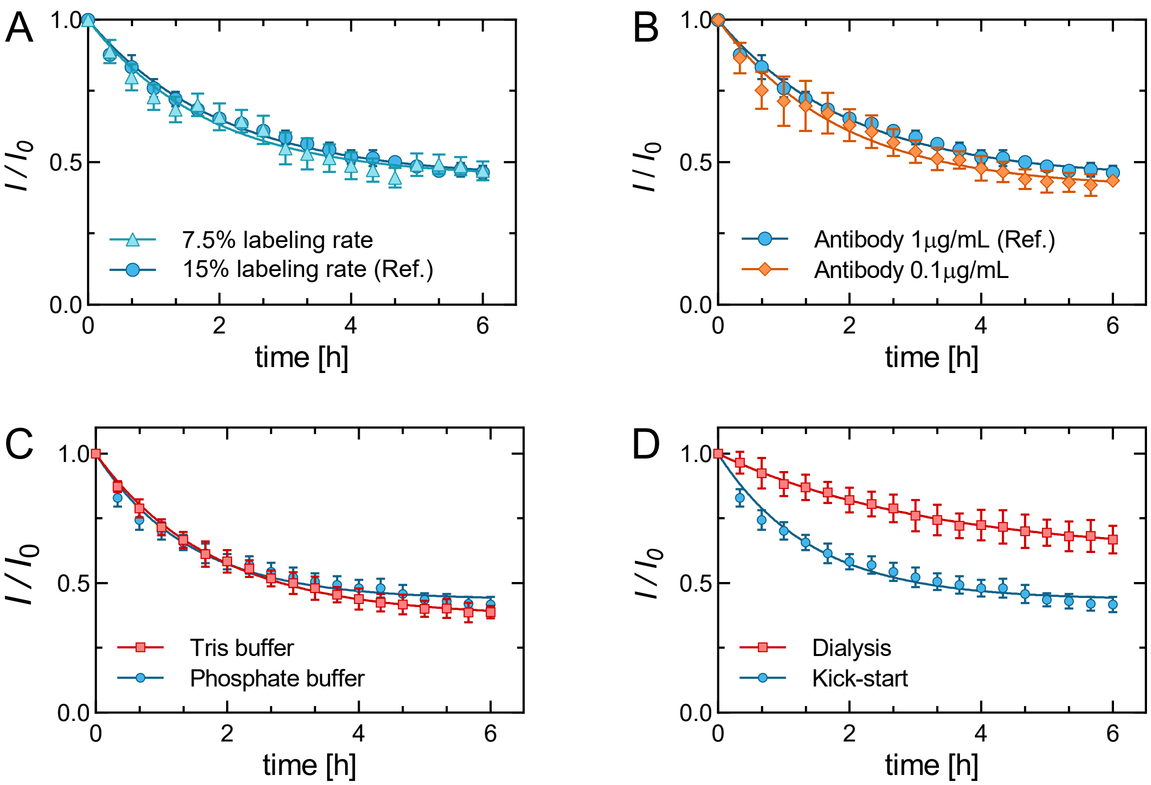


**Fig. S9.** Controls for the quantification of vimentin dissociation *in situ*. Impact of labeling fraction, concentration of antibodies, buffer composition, and method of filament assembly. (A) The fraction of fluorescent labeling does not impact filament dissociation. Time evolution of the mean fluorescence intensity normalized by the value at time 0 for vimentin assembled with an AF488- labeling fraction of 7.5% (light blue triangles) and 15% (blue circles), used as a reference in the paper. (B) The concentration of antibodies on the surface does not impact the dissociation curve, indicating that the level of tetramers trapped on the antibodies after dissociation is negligible. Time evolution of the mean fluorescence intensity normalized by the value at time 0 for vimentin assembled attached to a surface coated with a concentration of 1 µg/mL of antibodies used as a reference in the paper (blue circles) and with 0.1 µg/mL antibodies (orange diamonds). (C) Tris and phosphate buffer give the same dissociation curve. Time evolution of the mean fluorescence intensity normalized by the value at time 0 for vimentin assembled in Tris buffer (red squares) and phosphate buffer (blue circles) used as a reference in this paper. (D) Time evolution of the mean fluorescence intensity normalized by the value at time 0. Dialysed filaments (red squares) dissociate more slowly than filaments assembled by kick-start (blue circles). Each data point in (A-D) represents the average of $\sim$50 filaments per time point and the error bars are the standard deviations of the mean over 3 experiments. Solid line: fit using equation 6 of the Supporting Information, section: theoretical modeling.

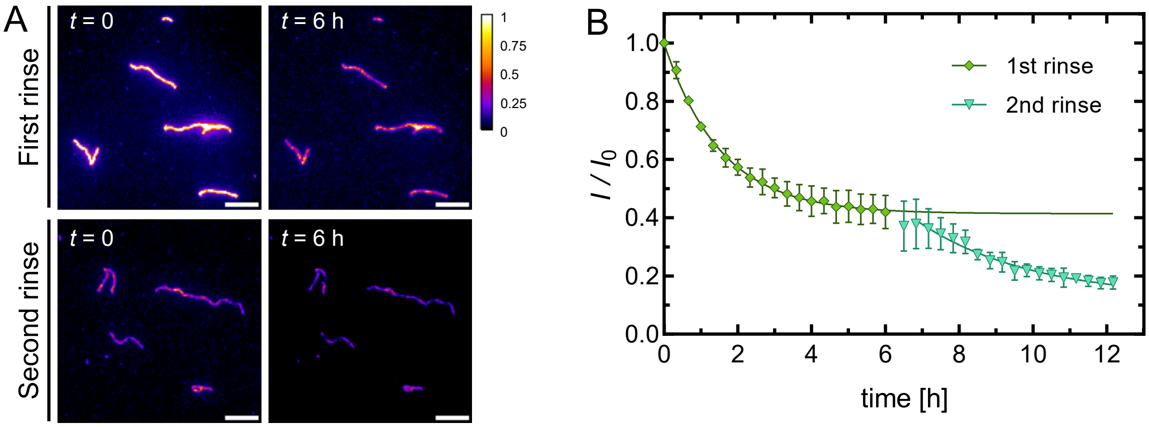


**Fig. S10.** In the *in situ* dissociation assay, successive rinsing of the observation chamber leads to two equilibrium levels. (A) Representative fluorescence images of filaments attached to the surface of the flow chamber at time 0, 6h after the first rinsing of the flow chamber and 6h after the second rinsing of the flow chamber. All images were acquired in similar conditions and are displayed with the same contrast (see the common color bar). Sale bar: 5 $\mu$m. (B) Time evolution of the mean fluorescence intensity normalized by the value at time 0, when the chamber is first rinsed. Each data point represents the average of $N = 3$ independent replicates ($\sim$50 filaments per condition and replicate) and the error bars are standard deviations of the mean over 3 replicates. Solid line: fit using equation 6 of the Supporting information, section: Theoretical modeling.

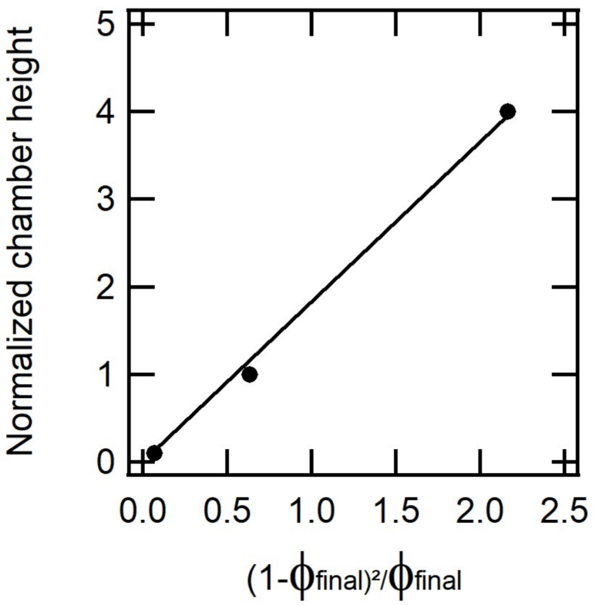


**Fig. S11.** Dependence of the immobile subunit fraction with the chamber's height. Eq. 14 of the theoretical modeling (Supporting Information) predicts that there is a linear relationship between the chamber height and $(1-\phi_{\mathrm{final}})^2/\phi_{\mathrm{final}}$, where $\phi_{\mathrm{final}}$ is the value of the parameter $\phi$, the fraction of binding sites that are occupied by a subunit, at equilibrium. Graph of the experimental chamber height normalized by the standard height $(H)$ corresponding to one layer of parafilm, as the function of the $(1-\phi_{\mathrm{final}})^2/\phi_{\mathrm{final}}$, where the values of $\phi_{\mathrm{final}}$ are obtained from the fit of the time curves in Fig. 3C in the three conditions: $0.1H$, $1H$ and $4H$ and corresponds to the values at equilibrium of $I/I_0$. The experimental points are well fitted by a line, confirming the linear relationship.

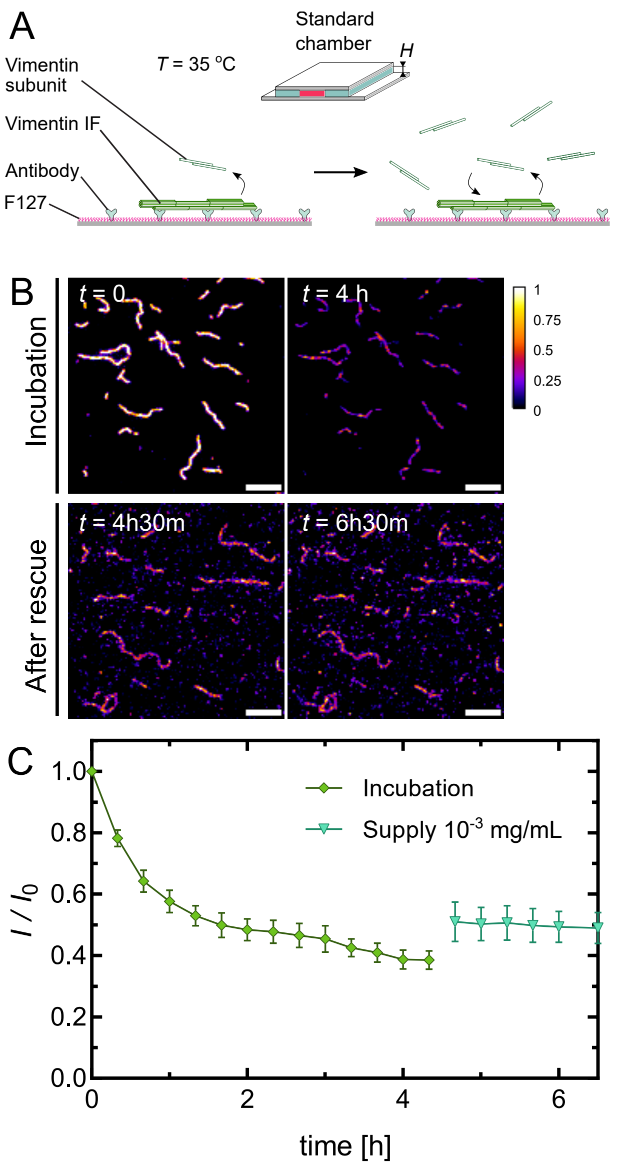


**Fig. S12.** Supply of $10^{-3}$ mg/mL tetramers to damaged filaments attached to the surface. (A) Schematic of the TIRF experimental setup. First, pre-assembled filaments were immobilized on the substrate for a duration of $\sim$4 h with the chamber being rinsed to remove initial free subunits. After incubation, we flowed in $10^{-3}$ mg/mL tetramers in the assembly buffer and continued the incubation. (B) Fluorescence images of filaments during the incubation step and after rescuing the damaged filaments with new tetramers. (C) Time evolution of mean fluorescence intensity normalized by the value at time 0 of filaments on the substrate during the incubation step and after rescue by supplying $10^{-3}$ mg/mL new tetramers. Each data point represents an average over 50 filaments and the error bars are standard deviations over 50 filaments.

### Supporting Information Text

### Methods and Materials

**Vimentin purification.** We purified recombinant wild-type human vimentin following Herrmann et al. 2004 (1). In brief, we transferred the vimentin construct in TG1 E. coli bacteria (Sigma-Aldrich) and started the induction when the OD reached 1.2. The induced bacteria were cultured in Terrific Broth medium overnight at 37 °C. Then, we collected the bacterial cells by centrifuging the culture medium, and lysed them with lysozyme, together with DNase (Roche), RNase (Roche), and protease inhibitors (Pefabloc and PMSF) in a 50 mM TRIS buffer pH 8. We washed the released inclusion bodies 5 times in the presence of DTT, and protease inhibitors: first with 200mM NaCl, 1% sodium desoxycholate, 1% NP40, 20mM TRIS pH 7.5, 2mM EDTA; second with 10mM TRIS pH 8, 0.5% Triton X 100; third with 10 mM TRIS pH 8, 0.5% Triton X 100 and 1.5M KCL; fourth with 10 mM TRIS pH 8, 0.5% Triton X 100; and finally with 10 mM TRIS, 0.1 mM EDTA pH 8 and 20μL DTT. Washing includes successive steps of incubation on ice for 20 min, centrifugation at 8000 rpm for 10 min at 4 °C, and resuspension using a cooled douncer. After washing, we resuspended the inclusion bodies in a denaturing buffer (8 M urea, 5 mM Tris pH 7.5, 1 mM EDTA, 1 mM DTT, 1 % PMSF) and centrifuged them at high speed (100000 $\times g$) for 1 h. Upon collecting the supernatant, we conducted vimentin purification by two sequential steps of exchange chromatography, using first an anionic (DEAE Sepharose, GE Healthcare) and then a cationic (CM Sepharose, GE Healthcare) column in a buffer of pH 7.5 containing 8 M urea, 5 mM TRIS, 1 mM EDTA, and 0.1 mM EGTA. We collected the vimentin protein in 2 mL Eppendorf tubes and monitored the protein abundance by Bradford assay. Only the most concentrated fractions were selected and pooled together. We stored the vimentin at -80 °C with an additional 10 mM methylamine hydrochloride solution.

For experiments, about 100 μL denatured vimentin aliquots were transferred to dialysis tubing (Servapor, molecular weight cut off at 12 kDa) and vimentin was renatured by stepwise dialysis from 8 M, 6 M, 4 M, 2 M, 1 M, 0 M urea into sodium phosphate buffer (pH 7.0, 2.5 mM sodium phosphate, 1 mM DTT) at room temperature with at least 15 minutes for each step. The final dialysis step was performed overnight at 4 °C in 2 L of the sodium phosphate buffer. The vimentin concentration was determined by UV-VIS absorbance measurements using a molecular weight of 53.65 kDa and extinction coefficient at 280 nm of 22,450 $M^{-1}cm^{-1}$ (2).

**Vimentin labeling.** We labeled vimentin proteins by forming a maleimide-thiol covalent bond between an Alexa Fluor 488 (or Alexa Fluor 555 and Alexa Fluor 647) maleimide dye and the cysteine-328 on vimentin (3, 4). In short, we dialyzed vimentin stored in 8 M urea for 3 h in a labeling buffer (50 mM sodium phosphate, pH 7, 5 M urea). The fluorescent dye (AF-488 C5 maleimide or AF-555 C2 maleimide or AF-647 C2 maleimide, ThermoFisher) resuspended in DMSO was pre-supplemented with a 5-molar excess. The solution was gently mixed for 1 h at room temperature, and the reaction was then quenched by the addition of DTT to 1 mM final. We used a Dye Removal Column (#22858, ThermoFisher) to remove excess dye. For storage, we dialyzed the vimentin collected after dye removal against a storage buffer (sodium phosphate 2.5 mM, pH 7, 8 M urea, 1 mM DTT) for 2 h at room temperature, then stored at -80 °C. For experiments, we renatured the labeled vimentin by stepwise dialysis from 8 M, 6 M, 4 M, 2 M, 1 M, 0 M urea to sodium phosphate buffer (pH 7.0, 2.5 mM sodium phosphate, 1 mM DTT). The labeled vimentin at 0 M urea was stored at 4 °C and used within 10 days.

**Vimentin filament assembly.** We assembled vimentin filaments for all the experiments using a standard kick-start assembly protocol, except specially notified. After dialysis in a buffer at 2.5 mM sodium phosphate pH 7.0, we added KCl to a solution of vimentin at the desired concentration and labeling ratio (typically 0.2 mg/mL with 15% labeling) with a final concentration of 100 mM KCl. Then, we placed the Eppendorf tube containing the vimentin mixture inside a water bath at 37 °C to initiate the assembly and incubated it for typically 2 hours. For the experiments where we needed to perform the assembly using the dialysis method, we put vimentin at the desired concentration and labeling ratio (typically 0.2 mg/mL with 15% labeling) into a piece of dialysis tubing (Servapor, molecular weight cut off at 12 kDa) in a 2 L beaker containing the assembly buffer (2.5 mM sodium phosphate, pH 7, 100 mM KCl) kept at 37 °C. KCl slowly diffused through the dialysis membrane and induced the assembly of the filaments. We typically assembled filaments for 12h at 37 °C.

**Mixing filaments.** We prepared two separate sets of unlabeled and AF-488-labeled vimentin (15% labeling fraction) filaments, both at 0.2 mg/mL. We assembled them simultaneously for 2 h at 37 °C in the assembly buffer (2.5 mM sodium phosphate, pH 7, 100 mM KCl). Then, we mixed the two sets of filaments at ratio of 1:1 or 9:1 and further incubated them at 37 °C for up to 24 h. We previously showed that pipetting does not break the filaments (4). At different time points during the experiment ($t = 0, 2, 4, ..., 24$ h), we took a small volume (10 μL) from the mixing solution and then immediately added an equal volume (10 μ L) of glutaraldehyde 0. 5% in the assembly buffer to fix the sample for later fluorescence imaging. Fixed filaments were diluted 100x in assembly buffer for further imaging. 4 μL of diluted filaments were placed between two glass coverslips and the sample was imaged with a standard epifluorescence microscope using a 60X objective, illumination with a LED lamp, and a digital CMOS camera. We verified that filament fixation did not affect the measurement of mean filament intensity or filament length (Fig. S1B). Furthermore, the imaging method, which involved placing a few microliters of fixed, diluted filaments between two non-passivated coverslips to which they adhered, did not alter the filament length distribution (Fig. S1C).

**Determination of vimentin soluble concentration.** After 2h of assembly at 0.2 mg/mL at 37°C with the addition of 100 mM KCl, without and with 15% vimentin labeling with AF488 (500 μ L per condition), the filaments were centrifuged at 140 000 $\times g$ for 15 minutes. The supernatant was collected and concentrated 10 times using centrifugal filter units (Amicon ultra, cut off at 30 kDa). The pellet was resuspended in 500 μ L of assembly buffer. We then measured the concentration of soluble vimentin in the supernatant by quantification of the band intensity on SDS-PAGE gels and comparison with a series of vimentin solutions of known concentrations used for calibration.

**SDS-page Gel and gel quantification.** We used 10% Mini-PROTEAN TGX precast polyacrylamide gels with 10 wells (Biorad). We loaded 10 μL of proteins per well mixed with a ratio 1:3 with 4X Laemmli Buffer Sample (Biorad) provided with 10% DTT. We ran the gels for 1 h at 120 V with Tris/Glycine/SDS buffer (Biorad) and stained the gels with InstantBlue protein stain (Expedeon) for 1 h. The intensity of the bands observed on the SDS-PAGE gels was analyzed using the plugin "Gels" from Fiji (5). The concentration of the vimentin soluble pool was estimated by comparing the intensity of the 10x concentrated supernatant bands with a range of vimentin pools with known concentrations used for calibration.

**Dilution experiments.** We first assembled AF-488-labeled vimentin filaments with 15% labeling fraction at 0.2 mg/mL for 2 h at 37 °C in the assembly buffer. For the dilution experiments in solution without tetramers, we diluted the filaments 1:200 or 1:500 in the assembly buffer. Then, we incubated the diluted samples for up to 6 h at 37 °C. We fixed the sample by taking a small volume of 10 μL and adding an equal amount of glutaraldehyde 0.5% in the assembly buffer. For the dilution experiments in solution with tetramers, we first added tetramers at desired concentrations and labeling fraction to the assembly buffer (for example, we prepared $10^{-3}$ mg/mL AF-488-labeled tetramers with 15% labeling fraction in the assembly solution). Then, we diluted the pre-assembled filaments 1:200 or 1:500 in the assembly buffer containing the pre-mixed labeled tetramers. We incubated the diluted samples for up to 6 h at 37 °C. We also fixed the sample by taking a small volume of 10 μL and adding an equal amount of glutaraldehyde 0.5% in the assembly buffer. Non diluted filaments were diluted 100X after fixation 1:1 in glutarahadelhyde 0.5%. Fixed filaments were imaged as the section "Mixing filaments": 4 μL of diluted filaments were placed between two glass coverslips and the sample was imaged with a standard epi-fluorescence microscope using a 60X objective, illumination with a LED lamp and a digital CMOS camera.

**Quantification of the subunit exchange.** For experiments mixing unlabeled and labeled filaments (Fig. 1), we assessed the extent of subunit exchange by quantifying the pixel fluorescence intensities along about 100 filaments per experiment extracted using linescan of 2 pixel width. The total distribution of pixel intensities showed two peaks: one at low intensity corresponding to the unlabeled filaments that incorporated fluorescent subunits, and one at high intensity corresponding to labeled filaments that incorporated unlabeled vimentin subunits (Fig. 1C). We fitted the distribution with double Gaussian that gave two mean intensities of the peaks: $I_{\text{low}}$ and $I_{\text{high}}$. We fitted the intensity of the labeled filaments before mixing with a single Gaussian, which gave their initial mean intensity $I_0$ (Fig. 1C). We then normalized the two peak intensities $I_{\text{low}}$ and $I_{\text{high}}$ by $I_0$ to assess the level of subunit exchange. The strength of this quantification method is that it does not need to detect whether the pixels were initially in labeled or unlabeled segments. We did not cut the filaments in small segments labeled vs. unlabeled but instead analyzed globally the distribution of fluorescence intensity along their length that we then attributed to labeled and unlabeled segments. We assume that a decrease in the mean fluorescence intensity of the bright peak results from the dissociation of labeled subunits and the association of unlabeled subunits. However, we cannot rule out the fact that this intensity decrease may come from the annealing of sub-resolved segments (<200 nm) of unlabeled filaments to labeled filaments. We argue that the distribution of filament length after 2h of assembly at 37°C displayed a very small proportion of small filaments below 200 nm (4).

**Quantification of filament length and MEAN fluorescence intensity.** For dilution experiments (Fig. 2-3), we used Fiji and 'Ridge detection' plugin to semi-automatically detect filament shape and length from fluorescence images. In detail, we first performed background subtraction for each original 16-bit image and created a duplicate of these images in 8-bit. Then, we used 'Ridge detection' on the 8-bit images and obtained detected filaments as ROI masks. Next, we manually validated the correctness of the detection and applied the ROI masks on the 16-bit images to obtain the length and fluorescence intensity values of the detected filaments. Notably, for fluorescence intensity measurement, we obtained the mean intensity along the detected lines with a linewidth of 2 pixels and set the minimum length of filament detection at 4 pixels to have an accurate fluorescence intensity measurement with minimal effect on the boundaries.

**TIRF microscopy experiment for direct observation of subunit dissociation along immobilized filaments on substrate.** We followed a protocol of TIRF microscopy experiment from our previous work (4). In short, we first cleaned coverslips by washing and sonicating them in a series of Hellmanex, KOH, and ethanol solutions. The coverslips were then completely dried, and treated in a plasma cleaner for 5 min. Immediately after plasma treatment, the coverslips were soaked in dichlorodimethylsilane 0.05% in trichloroethylene and incubated at room temperature for 1 h. We then washed off the excess silane by sonication of the coverslips 3 times in methanol for 15 min each. The silanized coverslips were dried by compressed air and stored at 4 °C for no longer than 2 weeks.

The flow chamber was assembled by melting pieces of parafilm (at about 50 °C) in between two coverslips. For the standard chamber, we assembled two coverslips with one layer of parafilm resulting in a chamber height $H$ of 100 μm. For the $4H$ chamber, we assembled 4 layers of parafilm in between the coverslips. For the $0.1H$ chamber, we made a thin chamber which includes 2 coverslips (18×18 mm and 22×50 mm) sandwiching a droplet of 3 μL and resulting in a height of 10 μm. For experiments to observe subunit dissociation *in situ* in a standard chamber or a $4H$ chamber, we flowed antibodies of AF-488 diluted 1:1000 in the assembly buffer (2.5 mM sodium phosphate, pH 7, 100 mM KCl) into the chamber and incubated for

5 min. Next, we flushed in 200 µL of F127 1% in the assembly buffer to the channel and incubated for 30 min for surface passivation. We then washed off the passivation agent by flushing an extensive amount of the assembly buffer. Pre-assembled AF-488-labeled vimentin filaments (assembly of 0.2 mg/mL with 15% labeling fraction for 2 h at 37 °C) were diluted 1:100 in the assembly buffer, then flushed into the chamber and incubated for 5 min. We washed all the unbound filaments and vimentin subunits out of the chamber by flushing in the assembly buffer only. We sealed the chamber with Vitrex sealant wax and transferred the device to a pre-heated microscope stage and objective. The temperature of the heating stage and objective was set to 35 °C.

For experiments with the $0.1H$ chamber, as solutions cannot be directly injected, we first pipetted two drops of 1:1000-diluted AF-488-antibody solution (500 µL) on a parafilm sheet, then placed two silanized coverslips on top of each drop to incubate for 5 min. We next placed each coverslip on top of a drop of passivation agent F127 solution (500 µL) and incubated for 30 min. We washed off the excess F127 by putting each coverslip on top of 3 drops of assembly buffer consecutively (500 µL for each drop). Then, each coverslip was transferred to a drop of 1:100-diluted pre-assembled filaments and incubated for 5 min. The surface was washed in 3 consecutive drops of assembly buffer (500 µL each), gently dried with a kimwipe (Kimtech). We deposited 3 µL of assembly buffer on one of the coverslips and placed the coverslips against each other to flatten the droplet into a thin layer of 18mm×18mm×10 µm. Finally, we sealed the thin chamber with vacuum grease and transferred it to the pre-heated microscope stage for imaging. TIRF imaging was carried out on a Nikon Eclipse Ti inverted microscope, equipped with a 60× oil-immersion objective. Images were captured by a Kinetix sCMOS Camera (Photometrics). The TIRF laser illumination was obtained from the Ilas2 module (GATACA Systems). We captured images of filaments on substrate with low laser power and a frame interval of 20 min. Before each experiment, we performed a bleaching control by acquiring the same number of images as in a long-term experiment, but with a shorter frame interval of 2 s (Fig. S8).

**Single molecule experiments.** We performed a series of single-molecule experiments to obtain characteristic bleaching steps of labeled vimentin subunits and tetramers. We used the same flow chamber setup as the one used in the TIRF microscopy experiment. In this experiment, to have a homogenous population of labeled vimentin tetramers, we first mixed unlabeled vimentin with AF-488-labeled vimentin at a monomeric level (in a denaturing buffer with 8M urea, 2.5mM sodium phosphate, 1mM DTT) to a desired labeling fraction of 15% or 50%. Then, we performed a stepwise dialysis of the labeled vimentin mixture from 8 M, 6 M, 4 M, 2 M, 1 M, 0 M urea to sodium phosphate buffer (pH 7.0, 2.5 mM sodium phosphate, 1 mM DTT). The collected vimentin tetramers at 0 M urea were stored at 4 °C for up to 10 days. For experiments to probe the bleaching steps of vimentin subunit dissociated from vimentin filaments, we flowed pre-assembled vimentin filaments labeled with AF-488 with 15% labeling fraction (assembly of 0.2 mg/mL for 2 h at 37 °C, and diluted 1:100 in the assembly buffer) into the chamber and incubated for 5 min. The unbound filaments and soluble subunits were rinsed off the chamber by flushing an extensive amount of the assembly buffer. The chamber was then sealed, and incubated for 60 minutes to let the subunits dissociate. To achieve high resolution for this single-molecule experiment, we used the Zeiss Elyra super-resolution microscopy system, equipped with a 100x oil-immersion objective and an EMCCD camera (Edge 4.2 CLHS (PCO)). The microscope stage and objectives were contained inside a microscope cage incubator operating at 37 °C (Pecon). We first set the laser illumination at low-power mode and recorded the binding time of newly recruited subunits on the substrate (capture setting: 0.5% laser power, 500 ms exposure time, 2 s time intervals between frames, total duration 2 min). Next, we set the illumination to high-power mode and captured newly recruited subunits on the substrate photo-bleached (capture setting: 2% laser power, 50 ms exposure time, no intervals between frames, total duration 10 min).

For experiments to probe the beaching steps of vimentin tetramers labeled with 15% and 50% labeling fractions, we made a flow chamber with only silanized coverslips. The silane can recruit vimentin tetramers rapidly and securely on the glass surface. To avoid overcrowding the surface with tetramers, we prepared the labeled tetramers at 0.1 mg/mL, diluted them 1:1000 in sodium phosphate buffer without KCl (2.5 mM sodium phosphate, pH 7), and then flushed them into the chamber for 30 s. We gently removed the unbound tetramers by flowing in a sodium phosphate buffer without KCl. We sealed the chamber and started the bleaching experiment at 37 °C using the same Zeiss Elyra microscopy system. Images were captured with high power mode settings (2% laser power, 50 ms exposure time, no intervals between frames, total duration 10 min). The distribution of bleaching steps expected theoretically for tetramers with a 15 % labeling fraction is: 77 % of all labeled tetramers should have 1 fluorophore, 20 % should have 2 fluorophores, 2 % should have 3 fluorophores, and 0.1 % should have 4 fluorophores. The theoretical prediction for tetramers with 50 % labeling fractions is: 27 % with 1 step, 40 % with 2 steps, 27 % with 3 steps, and 6 % with 4 steps. The probability of having $N$ fluorophores for the labeled tetramers with an average labeling fraction, $a$, is given by: $P = \frac{C(N,4)a^N(1-a)^{4-N}}{\sum_{k=1}^{4} C(k,4)a^k(1-a)^{4-k}}$, where C(N,4) is the number of combinations of having N fluorophores among the 4 molecules forming the tetramer.

**2D dSTORM imaging.** Samples were prepared as follows. Solutions of 20% labeled vimentin with AF-647 (i) assembled for 1h at 0.2 mg/mL, then diluted 40x in assembly buffer supplemented with 0.5% glutaraldehyde, (ii) assembled for 2 seconds at 0.2 mg/mL for 1h, then diluted 40x in assembly buffer provided with 0.5% glutaraldehyde, (iii) unpolymerized tetramers at $5 \times 10^{-3}$ mg/mL and (iv) assembled at $5 \times 10^{-3}$ mg/mL for 6h, were incubated on 18 mm clean coverslips for 10 minutes at room temperature. Then the coverslips were incubated with a solution of 100 nm tetraspeck beads (ThermoFisher #T7279) diluted at 1/500 in the imaging buffer for 15 minutes. Then the coverslips were mounted on a magnetic sample holder (Live Cell Instrument, CM-B18-1), with 1 mL of imaging buffer, and closed with a lit (6). 2D-dSTORM images were performed as described previously (7). Images were acquired on a Elyra-7 Zeiss microscope, using the TIRF µ-HD mode which focuses the intensity of the lasers in the center of the field of view, a ×63 NA 1.46 objective, and a 1.6 optovar lens and a sCMOS camera (Edge 4.2 CLHS (PCO)). We worked with 40×40 $\mu m^2$ field of view, used 20 ms exposure time and acquired 20 000 frames at the maximal intensity of the 642-nm laser power to pump most of the Alexa Fluor 647 dyes to the dark state ( 2 kW/$cm^2$) as described previously. The imaging buffer was made of Tris-NaCl buffer (50 mM Tris, 100 mM NaCl, pH = 8) supplemented with 10% of glucose (100 mg/mL), 10 mM MEA, 0.5 mg/mL (75 U/mL) glucose oxidase and 40 µg/mL catalase (80 - 200 U/mL). Images were reconstructed using the Zen software with a pixel size of 10 nm and using the Gaussian display mode with an expansion factor of 1 PSF. We obtained localizations with a precision below 20 nm. The drift was corrected using the model-based method.

For the quantification of the nearest neighbor distances, we first selected all the dots on the dSTORM images using the Analyze particles plugin applied with an intensity threshold of intensity 5. The nearest neighbor distances were then calculated using the "ndd" plugin from FIJI.

**AFM imaging.** We used an atomic force microscope (AFM, Nanowizard 4 from JPK/Bruker) to image vimentin filaments and estimate their diameters by measuring AFM heights. Fixed vimentin filaments were left to adsorb on glass coverslips (Menzel Glazer) that had been previously cleaned using deionized water and ethanol. After drying the coverslips with a N2 stream, we immersed 30 µL of vimentin filament solution on the coverslips for 5 min. The glass surface was rinsed 3 times with assembly buffer. We then used the AFM to image the vimentin filaments in liquid using the Quantitative Imaging (QI) mode and BL-AC-240TS tips (nominal spring constant: 0.1 N/m) from Olympus. Typically, we collected square images of 350 nm × 350 nm over 150 ×150 pixels. QI images were processed using the Data Processing software of JPK/Bruker. The height maps were obtained from force curves for each pixel, by collecting the piezo height and subtracting the cantilever deflection from it to obtain the vertical tip position at the point of contact between the tip and the filament surface or between the tip and the glass surface. To measure the mean diameter of a filament, we calculated the mean of the maximum height values for each cross-section along the filament (∼ 80 cross-sections per filament). .

### Theoretical modeling

Here we derive the theoretical expressions for the kinetics of vimentin subunit exchange and the resulting filament fragmentation discussed in the main text. In Sec. A we derive the simplest subunit turnover equation and use it to model the subunit dynamics in contact with an infinite reservoir used to model the rescue experiment of Fig. 3 of the main text. In Sec. B, we consider a slightly more complicated model that keeps track of both labeled and unlabeled subunits, as in Fig. 1. Section C then considers subunit exchange in contact with a finite experimental chamber, which we use in Fig. 4. Finally, in Sec. C we derive the average time for a filament fragmentation event under the assumption that the removal of a critical number of subunits is the associated rate-limiting step.

**A. Model for the turnover of mobile subunits.** We consider the exchange of vimentin subunits between a filament and an infinite solution containing a concentration $c$ thereof. The filament comprises a fixed number of immobile subunits, each of which can anchor one mobile subunit. We denote by $\phi(t)$ the fraction of occupied binding site at time $t$ in contexts where some of the subunits are immobile. More generally, $\phi(t)$ denotes the ratio of the number of subunits incorporated into the filaments to the maximum value that this number can take. Assuming that mobile subunits bind with a reaction rate $k_{\mathrm{on}}$ and unbind with $k_{\mathrm{off}}$, this fraction evolves according to

$$\frac{\mathrm{d}\phi}{\mathrm{d}t} = k_{\mathrm{on}} c(1-\phi) - k_{\mathrm{off}}\phi, \qquad [1]$$

where the term first term of the right-hand-side reflects that only a fraction $(1-\phi)$ of immobile subunits are available for binding. Denoting by $\phi(t=0)=\phi_0$ the initial fraction of occupied immobile subunits, we find that $\phi$ evolves to a final fraction

$$\phi_\infty(c) = \frac{k_{\mathrm{on}} c}{k_{\mathrm{off}} + k_{\mathrm{on}} c} \qquad [2]$$

according to

$$\phi(t) = \phi_\infty + (\phi_0 - \phi_\infty)\exp\left(-\frac{k_{\mathrm{off}} t}{1-\phi_\infty}\right). \qquad [3]$$

Assuming the measured filament fluorescence intensity $I(t)$ is proportional to the total density of subunits (immobile and mobile) along a vimentin filament and that about half of the total subunits are mobile as observed in the experiments, *i.e.*, to $(1+\phi)/2$, the filament fluorescence relative to its final state reads

$$\frac{I(t)}{I_\infty} = \frac{1+\phi(t)}{1+\phi_\infty} = 1 + \left(\frac{1+\phi_0}{1+\phi_\infty} - 1\right)\exp\left(-\frac{k_{\mathrm{off}} t}{1-\phi_\infty}\right), \qquad [4]$$

where we use the characteristic relaxation rate $k_{\mathrm{off}}/(1-\phi_\infty) = k_{\mathrm{off}} + k_{\mathrm{on}} c$ and the dimensionless relative intensity $(1+\phi_0)/(1+\phi_\infty)$ as adjustable parameters in fitting Fig. 3G of the main text.

**B. Exchange of labeled and unlabeled mobile subunits.** We now consider a situation where the fraction $\phi$ of occupied immobile subunits is equilibrated with the surrounding solution, implying $\forall t\ \phi(t) = \phi_\infty(c)$. We however also assume that only a fixed fraction $\lambda$ of the subunits in solution are labeled, implying that the fraction $\phi_l(t)$ of immobile subunits that are bound to a *labeled* mobile subunit may vary over time. The evolution equation for this last quantity thus reads

$$\frac{\mathrm{d}\phi_l}{\mathrm{d}t} = k_{\mathrm{on}}\lambda c(1-\phi_\infty) - k_{\mathrm{off}}\phi_l, \qquad [5]$$

the first term of whose right-hand-side differs from Eq. (1) in two ways. First, the concentration of soluble tetramers that lead to an increase of $\phi_l$ is now $\lambda c$ as opposed to $c$. Second, the fraction of immobile subunits available for binding is now $1-\phi_\infty$ and is thus fixed over time.

In the high-intensity filaments discussed in the text, all subunits composing the filament are initially labeled, *i.e.*, $\phi_l(t=0) = \phi_\infty$. Solving Eq. (5) with this initial condition leads to

$$\frac{I_{\mathrm{high}}(t)}{I_0} = \frac{1}{1+\phi_\infty}\left[1+\lambda\phi_\infty + (1-\lambda)\phi_\infty e^{-k_{\mathrm{off}}t}\right] \underset{t\to\infty}{\longrightarrow} \frac{1+\lambda\phi_\infty}{1+\phi_\infty}. \qquad [6]$$

By contrast, the low-intensity filaments do not initially comprise any labeled subunits, *i.e.*, have $\phi_l(t=0)=0$. Moreover their immobile subunits are unlabeled, implying $I/I_0 = \phi_l/(1+\phi_\infty)$ and leading to

$$\frac{I_{\mathrm{low}}(t)}{I_0} = \frac{\lambda\phi_\infty}{1+\phi_\infty}(1-e^{-k_{\mathrm{off}}t}) \underset{t\to\infty}{\longrightarrow} \frac{\lambda\phi_\infty}{1+\phi_\infty}. \qquad [7]$$

We note that these two expressions relax to their asymptotic values at rate $k_{\mathrm{off}}$ different than Eq. (4), which results from the different availabilities of the unbound immobile subunits in the two cases. We use Eqs. Eq. (6) and Eq. (7) to fit Fig. 1D in the main text (with $\lambda = 1/2$ and $\phi_\infty \simeq 1$ consistent with the observation that the mobile subunit binding sites are almost all occupied at equilibrium) and further compare its prediction to the $\lambda = 1/10$ data.

**C. Dilution into a finite-volume chamber.** We now consider a setup with a finite volume, *i.e.*, one where we cannot neglect the increase of the concentration of the pool of mobile subunits in solution as these subunits unbind from the filaments. This formalism is required to interpret the dilution experiments reported in Fig. 4 of the main text. We denote by $\rho$ the number of binding sites available to the mobile subunits per unit volume; $\phi$ is the fraction of these binding sites that are actually occupied by a subunit. This density may be computed as the product of the number of such sites per unit of filament length by the total length of the filaments in a certain observation volume, divided by this volume (which is itself proportional to the thickness of the chamber in Fig. 4B of the main text). We neglect any significant change in filament lengths throughout our experiments and thus treat $\rho$ as constant. Unlike in Sec. B, in the following, we do not consider different levels of subunit labeling.

As the amount of bound mobile subunits varies over time, subunits may be released into or taken up from the solution. As a result, upon an infinitesimal change $\mathrm{d}\phi$ of the fraction of occupied subunit binding sites, the concentration $c$ of mobile subunits in the solution changes by $\mathrm{d}c = -\rho\,\mathrm{d}\phi$. Following Fig. 4, we assume an initial condition where $c = 0$ and

$$\phi(t=0) = \phi_\infty(c_{\mathrm{inc}}), \qquad [8]$$

where $c_{\mathrm{inc}}$ is the vimentin concentration at which the filaments are incubated before their introduction into the flow chamber. This allows us to integrate the infinitesimal variation equation for $c$ and find $c = \rho[\phi_\infty(c_{\mathrm{inc}}) - \phi(t)]$.

Following the results of the previous section, we assume

$$\phi_\infty(c_{\mathrm{inc}}) \simeq 1. \qquad [9]$$

Inserting into Eq. (1) yields

$$\frac{\mathrm{d}\phi}{\mathrm{d}t} = k_{\mathrm{on}}\rho(1-\phi)^2 - k_{\mathrm{off}}\phi, \qquad [10]$$

which we combine with Eqs. (8-9) to find

$$\phi(t) = \frac{1+\alpha e^{-\tau}}{\alpha + e^{-\tau}}, \qquad [11]$$

where we have defined

$$\alpha = 1 + \frac{k_{\mathrm{off}}}{2k_{\mathrm{on}}\rho} + \sqrt{\frac{k_{\mathrm{off}}}{k_{\mathrm{on}}\rho} + \left(\frac{k_{\mathrm{off}}}{2k_{\mathrm{on}}\rho}\right)^2} \quad \text{and} \quad \tau = \left(4k_{\mathrm{off}}k_{\mathrm{on}}\rho + k_{\mathrm{off}}^2\right)^{1/2} t. \qquad [12]$$

In the experiment of Fig. 4C and E it appears that the fluorescence levels of the filaments can fall below 50% of their initial values, suggesting that in these specific experiments where the filaments are bound to the surface, all subunits are actually mobile. We model this observation by writing that unlike in the bulk experiments described in the last two sections, here $I(t)$ is proportional to $\phi(t)$. This implies

$$\frac{I(t)}{I_0} = \frac{1+\alpha e^{-\tau}}{\alpha + e^{-\tau}}. \qquad [13]$$

which we fit to the fluorescence curves of Fig. 4C and E to obtain the values $k_{\text{on}} = 5 \times 10^{-8}\,\text{M}^{-1}\cdot\text{h}^{-1}$ and $k_{\text{off}} = 2 \times 10^{-1}\,\text{h}^{-1}$. In addition to this change in the proportion of mobile subunits, the fit reveals that the surface binding modifies the exact values of these constants but not their order of magnitude (see discussion in the main text).

In the final equilibrium state of the system, Eq. (10) dictates that $k_{\text{on}}\rho(1-\phi_{\text{final}})^2 = k_{\text{off}}\phi_{\text{final}}$. As discussed in the first paragraph of this section, the volume of the chamber is moreover proportional to its height $H$, and therefore the binding site concentration is inversely proportional to H: $\rho \propto H^{-1}$. Combining these two conditions we find that

$$H \propto \frac{(1-\phi_{\text{final}})^2}{\phi_{\text{final}}}, \qquad [14]$$

which we experimentally confirm in Fig. S11.

**First-passage model of filament fragmentation.** We model the fragmentation of vimentin filaments by representing one filament cross-section as comprising at most $N$ mobile subunits bound to a core if immobile subunits. In practice, we use $N = 5$ as illustrated in Fig. 6 of the main text. Individual mobile subunits may stochastically leave the filament or rebind to it from the solution. We assume that the unbinding of the first subunit from a fully covered filament proceeds at a rate $k_{\text{off}}$. We furthermore note that the departure of a first subunit may locally weaken the filament, thus facilitating the departure of subsequent ones. We thus assume that the departure of subsequent subunits happens at a different rate, which we denote by $\alpha k_{\text{off}}$. We assume a single subunit rebinding rate $k_{\text{on}}c$, which is in particular valid at least in the case where rebinding is entirely diffusion-limited. We assume that the filament instantaneously fragments if $R$ of the $N$ subunits are missing, and seek to compute the mean of the time interval required for this to happen.

The simplest version of our model, which we discuss in the results section of the main text, considers $\alpha = 1$, *i.e.*, disregards this facilitation mechanism. In the more sophisticated version tackled in the discussion, $\alpha$ is larger than one and can be written as

$$\alpha = e^{\Delta\epsilon/k_BT}, \qquad [15]$$

where $k_BT$ is the thermal energy and $\Delta\epsilon$ may be interpreted as the difference between the binding energy of a subunit in a full filament cross-section *vs.* in a weakened, partially empty cross-section.

We denote by $n \in [0, N]$ the number of unoccupied sites in the cross-section, and consider the stochastic transitions between states characterized by different values of $n$. We denote by $g_n$ the transition rate from state $n$ to state $n+1$, and by $r_n$ the transition from state $n$ to state $n-1$. Starting from an initial condition $n = 0$, we compute the average mean first passage time $\langle\tau\rangle$ to the state $n = R$. According to Ref. (8), this quantity is given by

$$\langle\tau\rangle = \sum_{\nu=0}^{R-1}\sum_{\mu=0}^{\nu} \frac{r_\nu r_{\nu-1}\cdots r_{\mu+1}}{g_\nu g_{\nu-1}\cdots g_\mu}. \qquad [16]$$

The model described above dictates $g_0 = Nk_{\text{off}}$, $g_{n>0} = (N-n)\alpha k_{\text{off}}$ and $r_n = nk_{\text{on}}c$. In these expressions, the combinatorial prefactors stem from the fact that a system with $N-n$ mobile subunits loses a subunit $N-n$ times faster than a system with only one, and similarly for gaining subunits. Inserting these expressions into Eq. (16) yields

$$k_{\text{off}} \times \langle\tau\rangle = \frac{1}{\alpha}\sum_{\nu=0}^{R-1} \frac{(\alpha x)^{-\nu}}{(N-\nu)\binom{N}{\nu}}\left[\alpha - 1 + \sum_{\mu=0}^{\nu}\binom{N}{\mu}(\alpha x)^\mu\right], \qquad [17]$$

where we have used the notation $x = k_{\text{off}}/k_{\text{on}}c$.

In the case without facilitation ($\alpha = 1$), we combine the numerical values $k_{\text{off}} = 0.2\,\text{h}^{-1}$ and $k_{\text{on}} = 5 \times 10^8\,\text{M}^{-1}\text{h}^{-1}$ with a concentration $c = 100\,\text{nM}$. We find that for $R = 2$, *i.e.*, in the case where a filament breaks as soon as two mobile subunits are removed from the same cross-section, we have $\langle\tau\rangle \simeq 65\,\text{h}$, whose order-of-magnitude is on par with the fragmentation time of 18 h reported in Ref. (4). As a point of comparison, within this model values of $R = 1$ and $R = 3$ would imply $\langle\tau\rangle \simeq 1\,\text{h}$ and $\langle\tau\rangle \simeq 1.1 \times 10^4\,\text{h}$, respectively. To explore the effects of facilitation in more detail, we assume that $\langle\tau\rangle$ is equal to this measured value, and use $\alpha$, or equivalently $\Delta\epsilon$, as an adjustable parameter. Assuming $R = 3$, we find $\Delta\epsilon = 3.3k_BT$, while assuming $R = 4$ implies $\Delta\epsilon = 4.2k_BT$ and $R = 5$ yields $\Delta\epsilon = 4.9k_BT$. These numerical values underpin our discussion of the model in the results and discussion sections of the main text.
